# Gaining Biological Insights through Supervised Data Visualization

**DOI:** 10.1101/2023.11.22.568384

**Authors:** Jake S. Rhodes, Adrien Aumon, Sacha Morin, Marc Girard, Catherine Larochelle, Elsa Brunet-Ratnasingham, Amélie Pagliuzza, Lorie Marchitto, Wei Zhang, Adele Cutler, Francois Grand’Maison, Anhong Zhou, Andrés Finzi, Nicolas Chomont, Daniel E. Kaufmann, Stephanie Zandee, Alexandre Prat, Guy Wolf, Kevin R. Moon

## Abstract

Dimensionality reduction-based data visualization is pivotal in comprehending complex biological data. The most common methods, such as PHATE, t-SNE, and UMAP, are unsupervised and therefore reflect the dominant structure in the data, which may be independent of expert-provided labels. Here we introduce a supervised data visualization method called RF-PHATE, which integrates expert knowledge for further exploration of the data. RF-PHATE leverages random forests to capture intricate feature-label relationships. Extracting information from the forest, RF-PHATE generates low-dimensional visualizations that highlight relevant data relationships while disregarding extraneous features. This approach scales to large datasets and applies to classification and regression. We illustrate RF-PHATE’s prowess through three case studies. In a multiple sclerosis study using longitudinal clinical and imaging data, RF-PHATE unveils a sub-group of patients with non-benign relapsing-remitting Multiple Sclerosis, demonstrating its aptitude for time-series data. In the context of Raman spectral data, RF-PHATE effectively showcases the impact of antioxidants on diesel exhaust-exposed lung cells, highlighting its proficiency in noisy environments. Furthermore, RF-PHATE aligns established geometric structures with COVID-19 patient outcomes, enriching interpretability in a hierarchical manner. RF-PHATE bridges expert insights and visualizations, promising knowledge generation. Its adaptability, scalability, and noise tolerance underscore its potential for widespread adoption.

## Introduction

Data visualization via dimensionality reduction (DR) is an important component of exploratory data analysis in biology, especially with the recent explosion of high dimensional, high throughput biological data. Popular DR methods for visualization such as PHATE [1], t-SNE [2], UMAP [3], and others [4–6] are *unsupervised* and do not incorporate auxiliary data labels. Thus these methods provide visualizations that preserve the dominant structure in the data that is not necessarily relevant to data labels. As such, these methods may focus on features that are not of interest to practitioners. In this paper, we present a novel, *supervised* dimensionality reduction method intended for biological visualization.

In many applications, we may have access to metadata or annotations, either manually given or inferred, that we want to integrate with the data. The popular unsupervised DR-based visualization methods described previously do not use this auxiliary information during any step of the embedding process. Often such annotations or labels are expert-derived and can provide valuable insights into relationships between variables and label information. Thus there is a need for *supervised* DR-based visualization methods that incorporate this information.

Many unsupervised DR methods begin by computing pairwise distances or dissimilarities between all points. Thus some of these methods have been extended to the supervised setting by incorporating data labels directly into the distance measure [7–11]; however, these approaches suffer from several limitations. First, these class-conditional dissimilarity functions are order non-preserving [12], which means a dissimilarity function, *f*, does not guarantee *f* (*d*(*x, y*)) *< f* (*d*(*x, z*)) when *d*(*x, y*) *< d*(*x, z*) for any distance function *d*. Thus, these dissimilarity functions disrupt natural notions of distance, causing distortions in the relationships between data points that affect the visualization. Second, we will show that these methods tend to exaggerate the separation between observations of different classes, providing misleading visualizations and making downstream tasks unreliable. Third, the class-conditional dissimilarities are not defined for continuous labels (i.e. regression) nor extendable to unlabeled points.

We address these limitations with a new supervised DR method that we call random forest (RF)-PHATE. RF-PHATE uses a random forest to learn the relationships between the features and the supervised labels. The information is then extracted from the random forest and embedded into low dimensions, e.g., for visualization. The resulting embedding reflects the relationships between points when focusing on the features that are relevant to the supervised task while ignoring irrelevant features. RF-PHATE is scalable to large datasets and can be applied to classification and regression problems.

We demonstrate the ability of RF-PHATE to meaningfully embed data in three case studies; a visual overview is provided by Fig. 1. The cases include a longitudinal study of multiple sclerosis (MS) using clinical and imaging patient data (Fig. 1a-d), Raman spectral data collected from antioxidant-treated cancerous lung cells (A549) in which we demonstrate that antioxidants diminish the severity of diesel exhaust exposure to lung cells (Fig. 1ef), and COVID-19 patient plasma data, where we show that RF-PHATE visualizations align known antibody profiles with clinically-established patient-outcome labels in COVID-19 cases (Fig. 1hi. In each study, we show qualitatively and quantitatively that RF-PHATE provides clearer insights than competing methods.

**Fig. 1:**
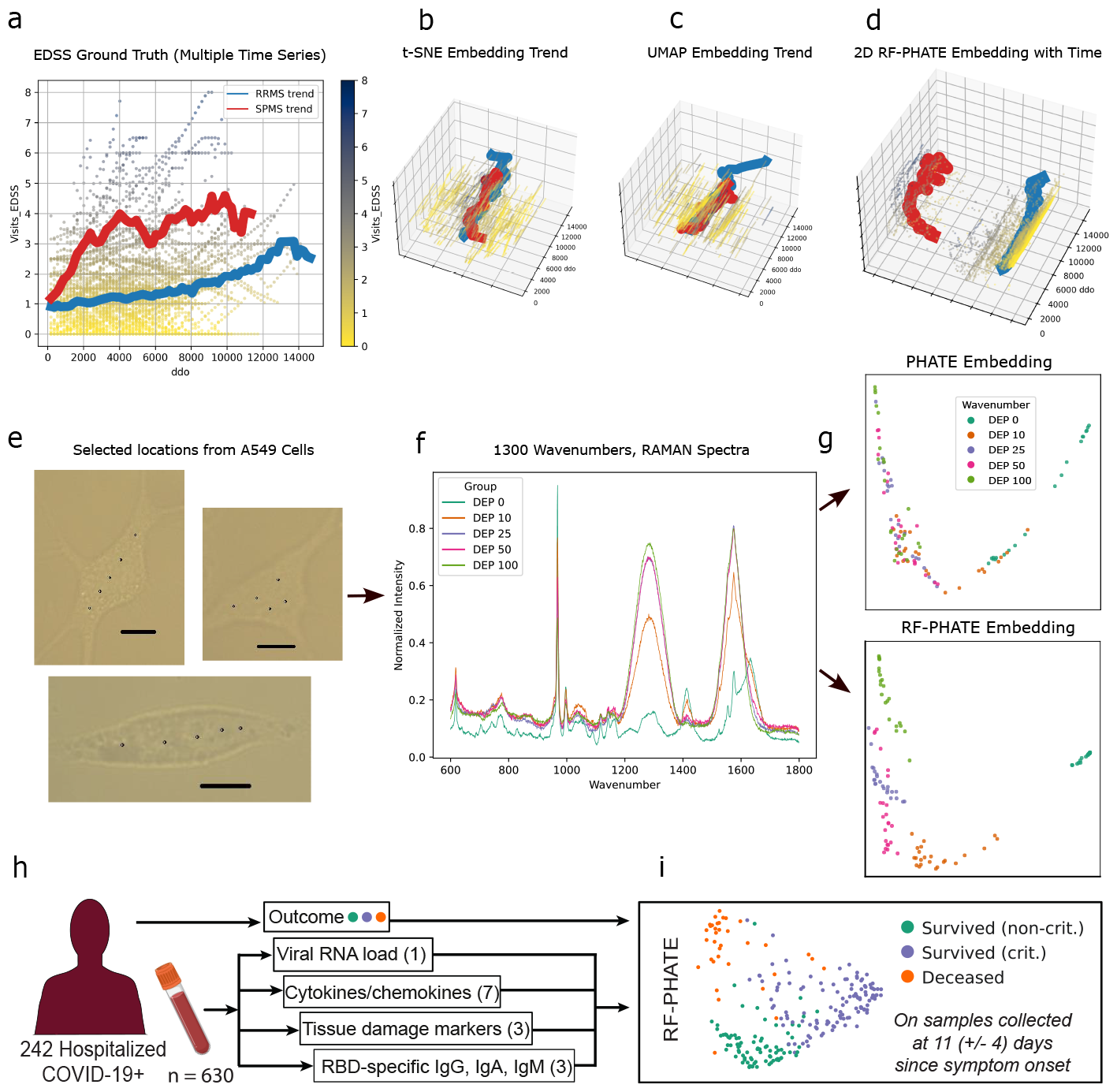
Overview of RF-PHATE and its ability to uncover latent, supervised structure. **a**) Multiple MS patient measurements of their expanded disability status scale (EDSS) over time, along with trend averages for RRMS and secondary progressive MS (SPMS) groups. **bcd**) 2-dimensional embeddings of the MS data with the trend averages (*t*-SNE, UMAP, and RF-PHATE shown here). Only RF-PHATE follows the same trend as the ground truth time series. **e**) A549 cell images and locations from which Raman spectra were recorded. **f**) Baseline-corrected Raman spectral values from diesel exhaust particle exposed A549 cells across five exposure levels. **g**) PHATE, and RF-PHATE embeddings are generated from the Raman spectra. The RF-PHATE embeddings show group separation without disrupting the overall trend. COVID-19 clinical data flow resulting in interpretable RF-PHATE embeddings (**i**).

We show that RF-PHATE visualizations highlight a sub-group of patients with non-benign relapsing-remitting MS (RRMS). In particular, our experiments show that two significant sub-groups of RRMS patients exhibit characteristics aligning with the traditional definition of benign MS (BMS) based on EDSS and disease duration. At the same time, other descriptors, such as disease worsening rate, annualized relapse rate, and treatment efficacy, distinguish our two RRMS sub-groups. A joint comparison with the SPMS group demonstrates that this distinction does not relate to RRMS patients being misclassified. Thus, our findings support the existence of benign and non-benign forms of RRMS defined as the combination of disease activity descriptors that better characterize the overall course of MS patients.

## Results

### RF-PHATE algorithm

RF-PHATE combines specialized RF proximities, known as RF geometry- and accuracy-preserving proximities (RF-GAP [14]), with the DR method PHATE to visualize the inherent structure of the features that are relevant to the supervised task while ignoring the irrelevant features. RF-PHATE has 5 main steps (Fig. 2):

**Fig. 2:**
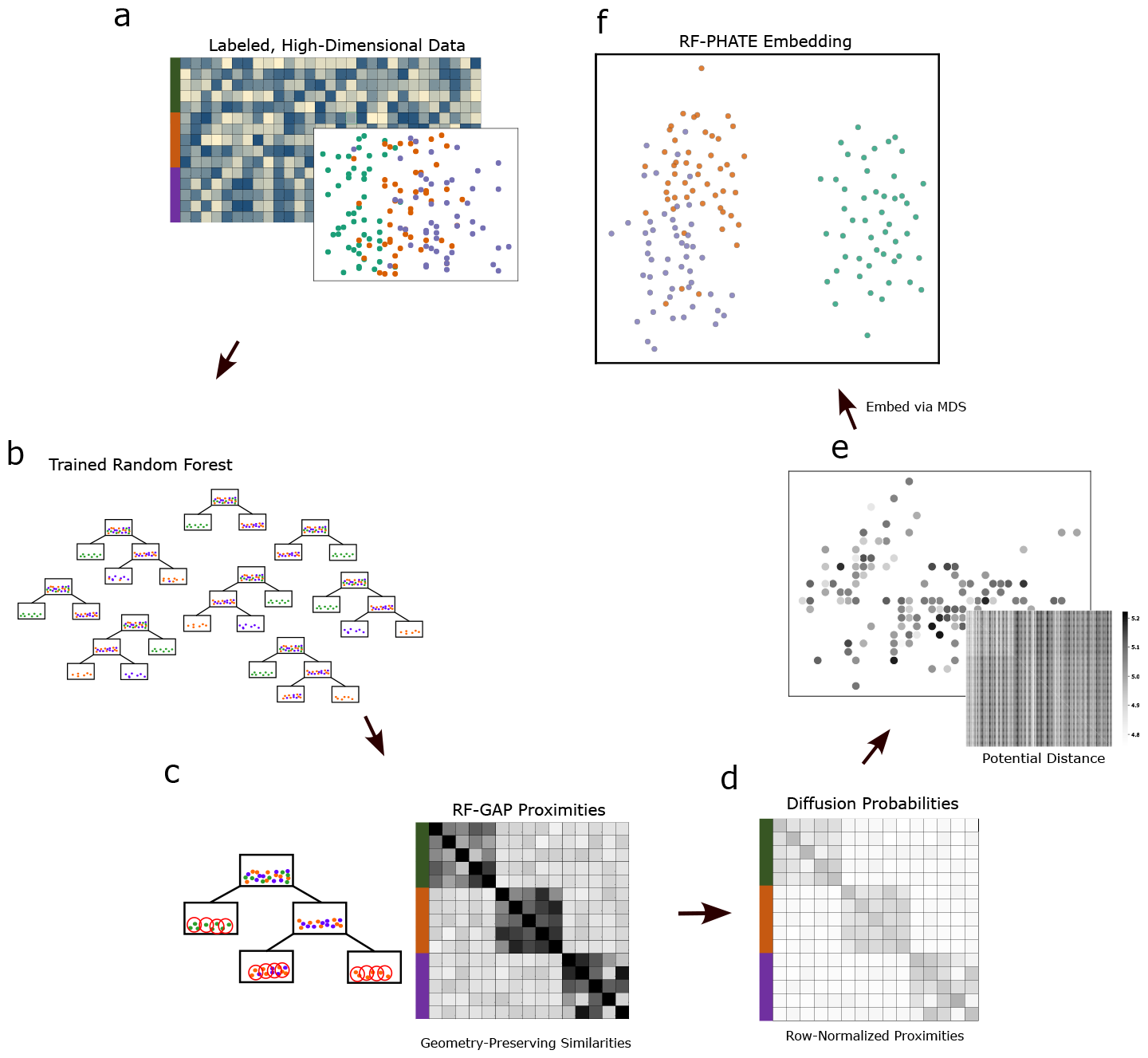
A visual representation of the RF-PHATE algorithm. **a**) Noisy, highdimensional data with labels is used to train a random forest [13] (**b**), from which local data similarities are computed using RF-GAP proximities [14] (**c**). The proximities are row-normalized, and damping [15] is applied to form the diffusion probabilities, which are stored in a Markov transition matrix (**d**). The global relationships are learned by diffusion, which is equivalent to simulating all possible random walks with time step *t* on the underlying Markov process. Mathematically, this is obtained by powering the transition matrix by *t*. A potential distance [1, 16] is then computed on the resulting diffused probabilities (**e**), which encodes both local and global relationships to which MDS [17] is applied to embed the data into low dimensions (**f**).

1. Train a random forest on the data to predict the labels (Methods and Fig. 2b).
2. Learn the local structure of the relevant variables using RF-GAP proximities [14] (Methods and Fig. 2c)
3. Learn the global structure of the relevant variables by diffusing the proximities (Methods and Fig. 2d)
4. Extract the local and global structure via the potential distances (Methods and Fig. 2e)
5. Visualize the structure by embedding the potential distances using metric multidimensional scaling (MDS) into 2 or 3 dimensions (Fig. 2f)

RF-PHATE first trains a random forest to predict the provided labels. Random forests are powerful supervised learning tools that provide accurate predictions with little to no tuning [18]. Additionally, random forests are suited for both classification and regression problems and can be formed between labeled and unlabeled points once the RF is trained. Thus they naturally extend to out-of-sample points.

By using random forest geometry-preserving similarity measures, we reveal a data structure that naturally incorporates known auxiliary information while keeping the benefits of the powerful random forest model. Unlike other RF proximity measures, RF-GAP proximities theoretically and empirically capture the inherent structure of the features that are relevant to the labeled data structure [14]. RF-GAP proximities do not artificiality distort distances, but form an alternative measure relative to label information. The use of RFGAP proximities overcomes the weaknesses of class-conditional dissimilarity measures. Thus, by using RF-GAP proximities as a local similarity measure, RF-PHATE is capable of learning a manifold structure that naturally incorporates label information while ignoring features that are irrelevant to the supervised task.

The RF-GAP proximities are computed between all pairs of points. The proximities serve as an adaptive kernel, *𝒫*, capable of encoding the local data structure while retaining important information for the supervised task. From *𝒫*, we form a row-stochastic diffusion operator, *P*, by normalizing each row in *𝒫*. Each entry in *P* represents local, single-step transitions between points. We then learn the global structure of the data by powering *P* by *t*-time steps, simulating all possible *t*-step random walks.

The selection for *t* is often overlooked in diffusion processes. A good choice of *t* should denoise and learn global structure without oversmoothing. We follow the construct of PHATE [1] and use Von Neumann entropy to identify the appropriate number of steps where denoising transitions to oversmoothing. The powered diffusion operator, *P*^*t*^, forms the basis for a potential distance that simultaneously retains local and global information which can be mapped to a lower-dimensional embedding. The potential distance, *D*_*t*_, is computed by log-transforming entries of *P*^*t*^ and then computing the pairwise distances between the log-transformed columns. Other works have shown this diffusion potential is capable of mapping out noisy and complex data in biological and other contexts [1, 16, 19]. The potentially complex, nonlinear structural relationships are finally mapped to a 2or 3-dimensional embedding, *G*, via metric MDS.

### RF-PHATE finds meaningful structure, filtering out irrelevant features

Our goal with RF-PHATE is to devise a method that generates a lowdimensional embedding that accounts for variable importance relative to its auxiliary (i.e., label) information. As such, noise variables (features that are not related to the auxiliary information) can and should be discredited or ignored in the embedding construction. To qualitatively show that RF-PHATE is capable of extracting meaningful visualizations relative to the supervised problem, we generated a high-dimensional (60), artificial tree dataset with a known structure (Methods and Fig. 3a) and added 500 uniformly-distributed noise variables. The labels are discrete and correspond to individual branches.

**Fig. 3:**
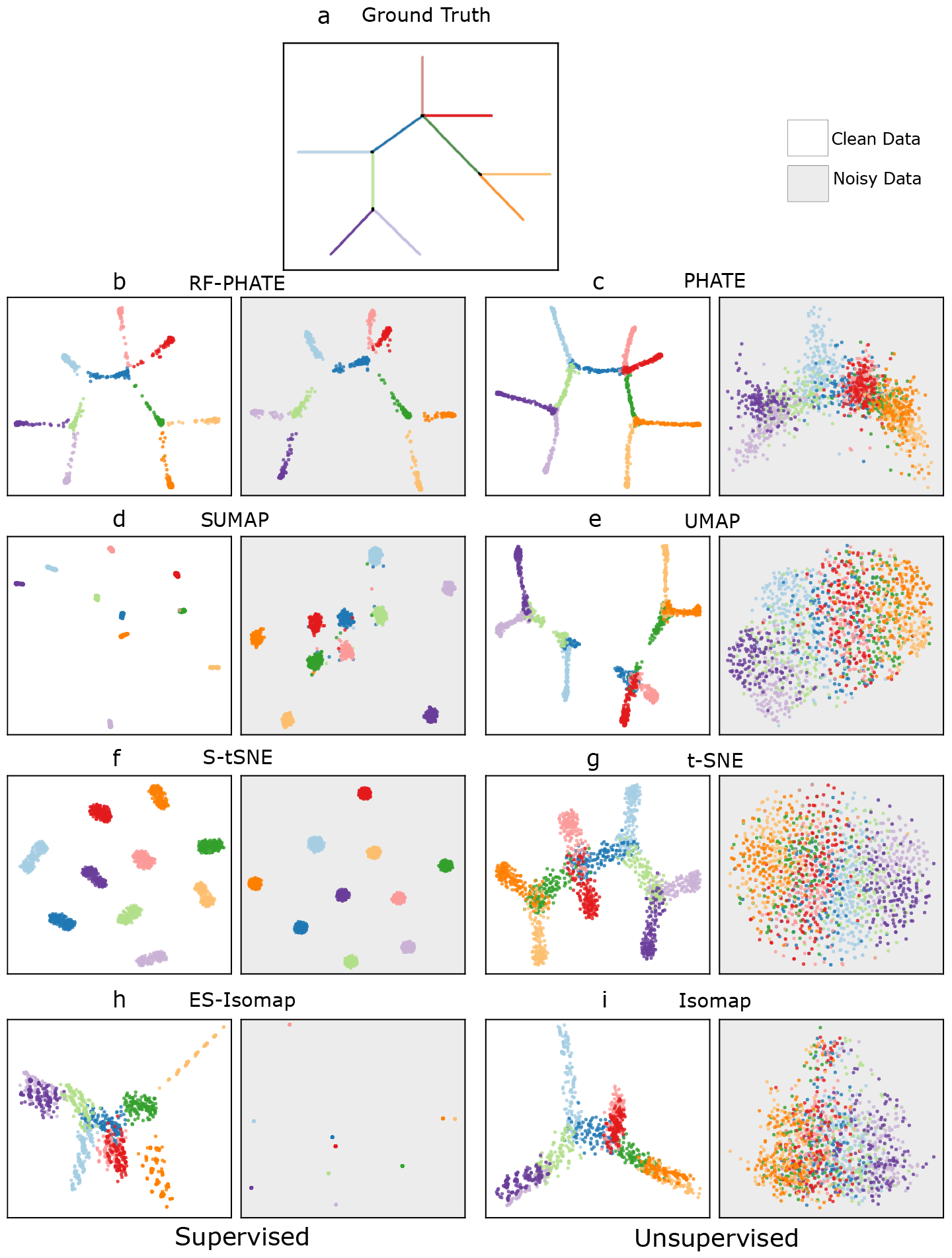
RF-PHATE is robust to noisy, irrelevant features. We generated an artificial tree structure (**a**), (the details of the tree’s construction can be found in Methods and [1]) and added 500 additional noise features (Methods). Without these noise variables (white background figures), RF-PHATE (**b**), PHATE (**c**), and to some degree t-SNE (**g**) accurately uncover the tree’s structure, while all other supervised (**d, f, h**) and unsupervised methods (**e, i**) fail. This shows that RF-PHATE works as well as unsupervised PHATE when the features that dominate the structure are also relevant for the supervised task. However, in the noise case (gray background), none of the unsupervised nor supervised methods recover the tree structure except for RF-PHATE.

Without the extra noise variables, unsupervised PHATE (Fig. 3c) provides an accurate depiction of the true tree structure. Unsupervised t-SNE also captures the structure fairly well in the sense that all ten branches are visible and appropriately connected to each other (Fig. 3g). However, UMAP and Isomap both introduce distortions (Fig. 3e,i). After including the noise variables, all of the unsupervised methods fail to represent the branching structure. In contrast, RF-PHATE accurately depicts the branch structure with or without noise (Fig. 3b) while the other supervised methods tend to separate the branches into distinct clusters (Fig. 3d,f,h). This provides a qualitative demonstration of RF-PHATE’s ability to represent the true structure of the data relative to the supervised task, in contrast with existing supervised and unsupervised DR methods.

Similar results were obtained on the Iris dataset (see Fig. S1 and Methods). In particular, RF-PHATE shows the true structure of the data with or without the noise variables, while other supervised methods introduce distortions by exaggerating class separation. These trends are corroborated numerically in the next section.

### Comparison of RF-PHATE to other methods

Here we quantitatively compare RF-PHATE with multiple unsupervised and supervised methods. Traditional performance metrics that evaluate an embedding’s goodness-of-fit are not suitable measures in our context since our focus is on features that are important for the supervised task. Such traditional methods include those based on pairwise distance preservation [5, 17], Procrustes analysis [20], or denoised manifold affinity preservation scores (DeMAP) [1].

As an alternative, we provide three new quantitative metrics to score the quality of low-dimensional embeddings in a supervised context (see Methods for details). 1) Low-dimensional group separation (LGS) determines the extent to which classes are distinguishable in low dimensions. 2) Local structure preservation score (LSP) measures the structure of local variable importance preservation in the embeddings, and 3) global structure preservation (GSP) relays information about global variable importance in low dimensions, where variable importance is measured with respect to the supervised task. A good supervised visualization method should perform well at all three metrics if it separates class without distorting the local and global structure present in features that are relevant for the supervised task.

In a wide comparison consisting of 27 publicly available datasets (see Methods and Table S4 for a description), we showed that RF-PHATE outperforms all unsupervised methods (10 total) using these metrics and on average outperforms existing supervised methods (14 total, including 5 RF methods that used the RF-GAP proximities as input). See Table 1. All metrics were performed on the two-dimensional embeddings. Thus based on these metrics, RF-PHATE outperforms other methods at learning an embedding that retains important information and structure relative to the supervised task in low dimensions.

**Table 1:** Compilation of results comparing embeddings across 24 DR methods, of which, 10 are unsupervised, 9 supervised, and 5 used RF-GAP proximities as an input. RF-PHATE outperformed all methods for both LGS and GSP and was bested only by S-tSNE for LSP. All scores were averaged across 27 publicly available datasets (Table S4) and 20 repetitions. Supervised methods (including RF methods) are marked with an asterisk.

| Method | LGS | LSP | GSP | Avg. Rank |
| --- | --- | --- | --- | --- |
| RF-PHATE* | **0.687 ± 0.01** | **0.780 ± 0.05** | **0.635 ± 0.05** | 1.33 |
| SUMAP* | **0.641 ± 0.04** | 0.754 ± 0.06 | **0.591 ± 0.12** | 2.33 |
| S-TSNE* [8] | 0.596 ± 0.02 | **0.790 ± 0.06** | 0.485 ± 0.08 | 3.67 |
| SNMF* [21] | 0.428 ± 0.06 | 0.647 ± 0.07 | 0.449 ± 0.08 | 7.33 |
| ES-LLE* [10] | 0.374 ± 0.02 | 0.561 ± 0.04 | 0.381 ± 0.07 | 9.67 |
| RF-UMAP* | 0.327 ± 0.03 | 0.716 ± 0.08 | 0.569 ± 0.10 | 9.67 |
| ES-ISOMAP* [9] | 0.485 ± 0.00 | 0.685 ± 0.03 | 0.379 ± 0.03 | 10.33 |
| RF-TSNE* | 0.324 ± 0.02 | 0.752 ± 0.08 | 0.570 ± 0.09 | 10.33 |
| NCA* [22] | 0.320 ± 0.00 | 0.557 ± 0.03 | 0.411 ± 0.03 | 11.00 |
| SPCA* [23] | 0.241 ± 0.00 | 0.494 ± 0.03 | 0.427 ± 0.03 | 11.33 |
| RF-ISOMAP* | 0.206 ± 0.03 | 0.662 ± 0.08 | 0.585 ± 0.09 | 14.33 |
| PLS-DA* | 0.184 ± 0.00 | 0.531 ± 0.03 | 0.205 ± 0.03 | 16.67 |
| T-SNE [2] | 0.183 ± 0.02 | 0.581 ± 0.07 | 0.406 ± 0.09 | 15.33 |
| KPCA | 0.149 ± 0.00 | 0.444 ± 0.04 | 0.335 ± 0.04 | 18.67 |
| PCA | 0.147 ± 0.00 | 0.439 ± 0.04 | 0.296 ± 0.04 | 19.33 |
| MDS [17] | 0.137 ± 0.01 | 0.471 ± 0.11 | 0.331 ± 0.05 | 19.00 |
| ISOMAP [5] | 0.131 ± 0.00 | 0.536 ± 0.03 | 0.340 ± 0.03 | 17.67 |
| UMAP [3] | 0.117 ± 0.02 | 0.517 ± 0.07 | 0.366 ± 0.08 | 19.00 |
| PHATE [1] | 0.107 ± 0.01 | 0.530 ± 0.04 | 0.441 ± 0.06 | 17.67 |
| KSPCA* [23] | 0.183 ± 0.00 | 0.581 ± 0.03 | 0.267 ± 0.03 | 16.67 |
| LAP. EIG. [6] | 0.085 ± 0.00 | 0.495 ± 0.03 | 0.426 ± 0.03 | 14.67 |
| RF-MDS* [18] | 0.070 ± 0.02 | 0.516 ± 0.10 | 0.378 ± 0.11 | 15.67 |
| DM [24] | 0.032 ± 0.03 | 0.437 ± 0.06 | 0.281 ± 0.09 | 18.67 |
| LLE [4] | -0.091 ± 0.01 | 0.544 ± 0.05 | 0.374 ± 0.06 | 21.33 |

### RF-PHATE outperforms other methods in visualizing MS follow-up data in supervised settings

In this study, we extended RF-PHATE to time-series data and identified key trends underlying the course of MS. We pioneered the use of MS patient timeseries data from disease onset by leveraging a unique MS database generated in the Centre de Recherche du Centre Hospitalier de l’Université de Montréal (CRCHUM), as described in Methods and Tables S1 and S2. Pairwise feature correlations can be found in Fig. S2. Diagnoses were retrospectively labeled by domain experts at the last follow-up as one of the two major forms of MS: RRMS or SPMS (secondary progressive MS). While these two forms are usually differentiated by a single progressive feature, namely the expanded disability status scale (EDSS [25, 26]), we explored other longitudinal descriptors such as functional system scores or MRI data to clinically and physiologically characterize RRMS and SPMS trajectories to potentially refine sub-types of RRMS and SPMS. Through RF-PHATE, we constructed a visualization of long-term MS courses that allows domain experts to easily retrieve known differences between MS classes (domain knowledge retrieval) and detect potential between- and within-class differences in terms of additional longitudinal features (knowledge discovery).

We used existing labeling to project our 16-dimensional MS follow-up database into 3 dimensions using RF-PHATE and compared it with 23 other unsupervised and supervised DR algorithms using the same three metrics as before (Table 2), although they were moderately adapted to better suit timeseries data (see Methods). We again showed that RF-PHATE outperforms other methods in embedding the original data while both separating the two classes and preserving the local and global structure with respect to the most important features of RRMS/SPMS classification.

**Table 2:** Performance scores of 24 different embedding methods on MS timeseries data using our three benchmark metrics LGS, LSP and GSP. Supervised embedding methods are marked by an asterisk. The numerical values are given as mean*±*standard deviation. We put the second-highest scores in bold, while the top scores are also underlined. Methods are sorted by their average rank across the three metrics (last column). RF-PHATE is the only embedding method ranked first (GSP) or second (LGS, LSP), making it the best overall method to embed our MS longitudinal database while separating RRMS/SPMS trends and preserving the local and global structure of the original data in terms of relevant features. While SNMF outperforms RF-PHATE in the LGS comparison, its almost perfect score is symptomatic of overly separated classes (Fig. S4). Moreover, although PLS-DA performs slightly better in the LSP comparison, it still does not effectively distinguish between the classes, as evidenced from its low LGS score and its embedding plot presented in Fig. S4.

| Method | LGS | LSP | GSP | Avg. Rank |
| --- | --- | --- | --- | --- |
| RF-PHATE* | **0.797 ± 0.01** | **0.756 ± 0.00** | <u>0.934 ± 0.00</u> | 1.67 |
| SPCA* | 0.471 ± 0.00 | 0.700 ± 0.00 | **0.903 ± 0.01** | 3.00 |
| SNMF* | <u>0.999 ± 0.00</u> | 0.647 ± 0.00 | 0.901 ± 0.01 | 3.67 |
| PLS-DA* | 0.435 ± 0.00 | <u>0.779 ± 0.00</u> | 0.882 ± 0.00 | 4.33 |
| RF-ISOMAP* | 0.251 ± 0.02 | 0.664 ± 0.05 | 0.900 ± 0.05 | 7.00 |
| KSPCA* | 0.304 ± 0.00 | 0.677 ± 0.00 | 0.856 ± 0.00 | 7.67 |
| RF-TSNE* | 0.192 ± 0.01 | 0.676 ± 0.04 | 0.885 ± 0.05 | 8.67 |
| MDS | 0.299 ± 0.00 | 0.610 ± 0.04 | 0.858 ± 0.01 | 9.00 |
| ES-ISOMAP* | 0.476 ± 0.00 | 0.421 ± 0.00 | 0.809 ± 0.00 | 9.67 |
| PCA | 0.349 ± 0.00 | 0.526 ± 0.00 | 0.641 ± 0.00 | 10.00 |
| RF-UMAP* | 0.215 ± 0.02 | 0.620 ± 0.07 | 0.846 ± 0.05 | 10.33 |
| ES-LLE* | 0.341 ± 0.23 | 0.489 ± 0.18 | 0.616 ± 0.29 | 11.33 |
| RF-MDS* | 0.090 ± 0.00 | 0.605 ± 0.03 | 0.897 ± 0.02 | 11.33 |
| NCA* | 0.326 ± 0.00 | 0.453 ± 0.00 | 0.351 ± 0.04 | 12.67 |
| KPCA | 0.130 ± 0.00 | 0.515 ± 0.00 | 0.488 ± 0.00 | 15.00 |
| ISOMAP | 0.176 ± 0.00 | 0.241 ± 0.00 | 0.662 ± 0.00 | 16.00 |
| SUMAP* | 0.198 ± 0.04 | 0.370 ± 0.14 | -0.104 ± 0.14 | 16.67 |
| S-TSNE* | 0.204 ± 0.04 | 0.262 ± 0.14 | -0.210 ± 0.07 | 17.67 |
| LLE | 0.234 ± 0.00 | -0.171 ± 0.00 | -0.062 ± 0.00 | 17.67 |
| LAP. EIG. | 0.020 ± 0.00 | 0.356 ± 0.00 | -0.129 ± 0.00 | 19.00 |
| DM | -0.677 ± 0.00 | 0.206 ± 0.00 | 0.294 ± 0.00 | 20.33 |
| UMAP | -0.068 ± 0.02 | 0.142 ± 0.10 | -0.457 ± 0.05 | 22.00 |
| PHATE | -0.029 ± 0.00 | -0.221 ± 0.01 | -0.342 ± 0.04 | 22.33 |
| T-SNE | -0.075 ± 0.01 | -0.115 ± 0.11 | -0.573 ± 0.06 | 23.00 |

Although supervised nonnegative matrix factorization (SNMF) [21] has a higher LGS score than RF-PHATE, its almost perfect score is symptomatic of overly separated classes. Indeed, SNMF fails to reach favorable performance in LSP, indicating a lack of structure preservation corroborated by its corresponding plot (Fig. S4) in which trajectories of RRMS and SPMS patients are compressed within their respective clusters, making it difficult to extract any meaningful patterns. RF-PHATE and other competing methods showing relatively good LSP and GSP scores, such as supervised principal component analysis (SPCA [23]) or partial least squares-discriminant analysis (PLS-DA), are able to preserve the EDSS structure from the ground truth plot (Fig. S4), with RRMS and SPMS grouped motions starting from similar regions, and SPMS quickly diverging over time towards higher EDSS values. Since EDSS has been shown to be the most important feature in the RRMS/SPMS timeseries classification (Table S5), this confirms the ability of these methods to embed patient trajectories while preserving the most important longitudinal structure of the original data.

However, unlike RF-PHATE, SPCA and PLS-DA fail to demonstrate class separability and establish between- and within-class characterizations. Indeed, they fall far behind RF-PHATE in the LGS comparison. This is reflected visually in Fig. S4, as they do not highlight the well-separated clusters of RRMS and SPMS patient time series. As for the other embedding plots, they exhibit comparable weaknesses, if not more pronounced ones (Fig. S4). Thus, we have shown, both quantitatively and qualitatively, that RF-PHATE is the best overall method to embed our longitudinal MS database in low dimensions while separating known MS types and preserving the most important longitudinal structure from the original space. This shows that RF-PHATE is a strong approach to visualize and compare typical MS trajectories in a supervised context.

### RF-PHATE provides insights into benign MS

Benign MS (BMS) is a sub-type of RRMS in which patients experience a mild course of disease that includes, for example, no severe relapses, complete recovery from relapses, or no progression outside of relapses. Given the heterogeneous nature of the disease, MS appears to encompass a sub-group of RRMS individuals who, even after numerous years, undergo minimal physical disability. While BMS is frequently observed in MS patients, there is a lack of consensus both in terms of defining this type of MS and acknowledging its existence [27, 28]. While the current standard definition of BMS is solely based on sustained EDSS scores below some threshold value [29], the practical significance of this entity in clinical practice remains uncertain [29]. This controversy arises from studies pointing out that symptoms such as fatigue, depression, cognitive dysfunction, and pain are not accounted for by the current definition of BMS despite their potentially great impact on patients’ well-being and work ability [30–34]. Hutchinson [35] stated that truly benign MS is rare and strongly discouraged the clinical use of the traditional definition because it may lead clinicians to become complacent or overly reassuring in their treatment plans. Yet, other studies found that basic EDSS-based definitions of BMS could be sufficient to characterize patient well-being and workability over time [36–38], despite variations in definitions across studies regarding EDSS score (0–4) and disease duration (5–15 years) [39].

In response to this controversy, several research groups have suggested alternative approaches to defining BMS. Recently, Morrow et al. [40] proposed an improved version of EDSS to consider cognitive decline during relapses. Ellenberger et al. [41] introduced a conservative definition (EDSS ≤ 1.0, absence from any disability, and the ability to work after 15 years of disease duration) to better reflect patients’ well-being in their everyday life. From a biomarker standpoint, Correale et al. [42] reported the potential of novel quantitative MRI techniques and further assessment of spinal cord abnormalities in detecting BMS and distinguishing it accurately from other forms of the disease. In this direction, Golan et al. [43] found that computerized cognitive scores were significantly associated with quantified MRI. Additionally, Niiranen et al. [44] identified grey matter volume as a significant measure for distinguishing between EDSS-based BMS and non-BMS cases. To summarize, there is still no general agreement for the definition for BMS, but focus has shifted toward an updated definition which covers a fuller spectrum of MS disease activity.

A powerful aspect of RF-PHATE is its ability to simultaneously consider other factors, such as the eight functional scores of EDSS and imaging data, to stratify the RRMS group into two sub-groups that could better define the distinction between benign and non-benign RRMS. Moreover, its capability to consider input features according to their importance to the RRMS/SPMS classification task means that the overall structure of the visualization is related to the long-term progression of the disease.

A detailed examination of the RF-PHATE embedding of the MS data uncovers a more severe and unstable sub-group of RRMS. The RF-PHATE plot (Fig. 4a) provides evidence for the presence of this sub-type with its capacity to distinguish between the RRMS and SPMS classes while maintaining the underlying structure. Indeed, our RRMS sub-typing (Fig. 4b) discovers 2 clusters of patient trajectories of large sizes (52 and 105 patients for the light blue and dark blue clusters, respectively), showing a significant spatial distinction. Visually, the light blue cluster exhibits RRMS trajectories characterized by high spatial variability, while the dark blue cluster gathers all of the stable RRMS trajectories. Since we have established in earlier experiments that RF-PHATE preserves the original structure, we are confident that this clear spatial differentiation directly relates to important differences within RRMS patient trajectories in terms of the input features.

**Fig. 4:**
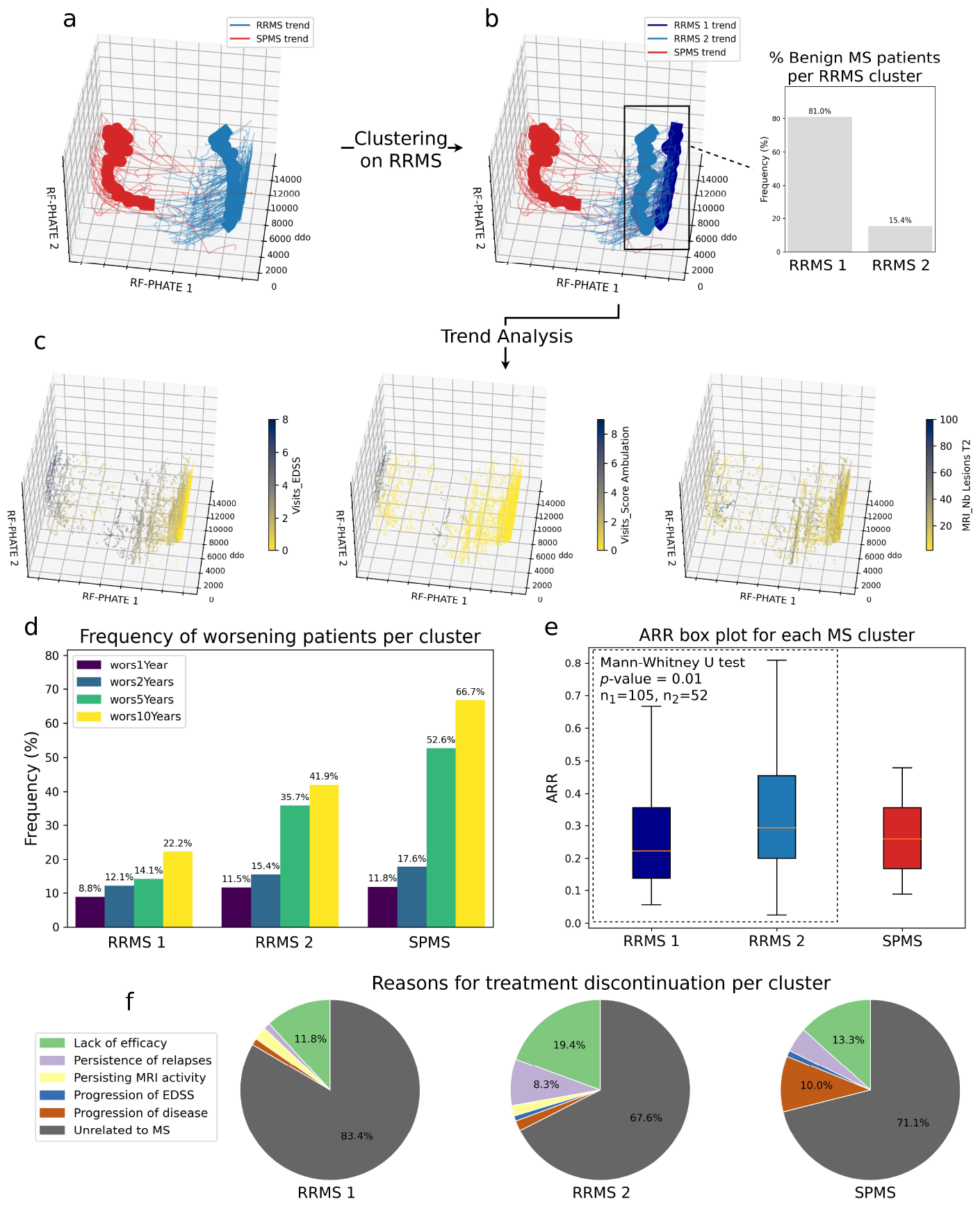
Overview of the exploratory analysis of MS follow-up data using RF-PHATE. (**a**) Three-dimensional RF-PHATE embedding plot of MS patient trajectories colored by the two existing RRMS/SPMS classes. (**b**) Three-dimensional RF-PHATE embedding plot of MS patient trajectories colored by SPMS along with two sub-groups of RRMS detected using 2-medoids and dynamic time warping (DTW, see Methods). Bold lines show the global motion, computed as the time-wise average coordinates of the respective groups. RF-PHATE unveils two sub-groups of RRMS patients with very different profiles. Indeed, the light blue cluster exhibits high spatio-temporal variability and diverges from the dark blue cluster which consists of stable trajectories concentrated in the right portion of the plot. Moreover, thanks to RF-PHATE’s ability to preserve the longitudinal structure of the data, the proximity observed between the RRMS 2 and SPMS clusters indicates similarities between these two groups of patient trajectories. (**c**) RF-PHATE plot of patient visits colored by 3 input features. (**d**) Frequency of patients worsening after *i* = 1, 2, 5, 10 years within each cluster. (**e**) Annualized relapse rate (ARR) distribution within each MS cluster. (**f**) Pie charts showing reasons for treatment discontinuation per cluster.

Fig. 4c highlights three specific features that contribute to this differentiation (see Fig. S5 for the whole input feature set). The bottom-left part of the RRMS cluster shows a notable increase in values for EDSS (Fig. 4c, left) and nearly all other clinical indicators (Fig. S5). This suggests that the two detected RRMS sub-groups make an allusion to two time-series clusters associated with benign and non-benign RRMS trajectories that are explained clinically. From a physiological perspective, the plots color-coded by MRI features do not clearly exhibit a distinct course toward a lower or higher number of brain lesions within each of the two RRMS clusters, except for a slight rise in the number of T2 lesions within the light blue cluster (Fig. 4c, right and Fig. S5). This implies that the overall clinical worsening of the light blue cluster in Fig. 4b is not necessarily linked to a progressive increase in the number of lesions over time, regardless of the brain region involved. However, this does not mean that there is no link between MRI data and clinical symptoms in the long run. Indeed, categories used in the McDonald criteria [45] do not seem refined enough to quantify meaningful radiological patterns. For instance, the McDonald Lesions T2 and McDonald Lesions Periventricular plots mostly consist of visits with scores 9+ and 3+, respectively, which fails to capture higher levels of quantification. Therefore, visually detecting physiological differences from our data is challenging, and further investigation is conducted in the next section.

At the same time, the light blue time-series cluster is similar to the neighboring SPMS cluster in terms of clinical and radiological features. This means that the light blue time-series cluster may also contain patients mislabelled as RRMS in place of SPMS. Indeed, SPMS patients could have been intentionally classified as RRMS to alleviate the financial and emotional burdens associated with being categorized as a more severe form of MS (“therapeutic mislabelling” [46]). It is also possible that SPMS was not diagnosed in RRMS patients due to the gradual nature of functional decline, which often goes unnoticed by both patients and doctors. This phenomenon, known as “silent progression” [47, 48], is observed in the RF-PHATE plots (Fig. 4c). While the second RRMS branch shares very similar features to the SPMS group, the latter generally displays high feature values earlier after disease onset (cf. Score Mental and Score Visual sub-plots of Fig. S5). Still, poor scores in Ambulation (Fig. 4c, center), Brainstem, and Cerebellar (Fig. S5) remain very specific to the SPMS cluster, which rather supports the existence of a distinct RRMS sub-group. Acknowledgment of this sub-group may have future implications on patient treatment, which is not currently optimized for this specific sub-group (see Fig. S6).

### Quantitative investigation supports the existence of a more severe RRMS sub-group

Using the previous visual observations as a starting point, we examined other accessible patient data to provide a rationale for the distinct behaviors exhibited by the three clusters of interest, namely RRMS 1 (dark blue in Fig. 4), RRMS 2 (light blue), and SPMS (red).

We wanted to verify whether the two RRMS sub-clusters were indeed associated with benign and non-benign MS courses. To do so, we first used a basic definition of BMS that combines the EDSS score with disease duration. That is, BMS was identified in a patient if and only if their EDSS score was 3 or lower for a disease duration of 20 years or more, or if the EDSS score was 2 or lower for a disease duration of 10 years or more. Consequently, out of the 105 patients in the dark blue cluster, 85 (81.0%) were assigned with BMS, whereas only 8 out of the 52 patients (15.4%) in the light blue cluster were assigned with BMS (Fig. 4b), which confirmed our previous statement. Another aspect of long-term clinical characterization regarding MS disease progression is the worsening of MS symptoms, as defined in Methods. Fig. 4d highlights a large increase in the frequency of worsening patients after 5 years since disease onset within the RRMS 2 (non-benign) cluster which is not apparent within the RRMS 1 (benign) cluster. Indeed, while the frequency of worsening patients within the RRMS 2 cluster remained below approximately 15% up to 2 years since disease onset, it escalated to 35.7% and 41.9% after 5 and 10 years, respectively. In contrast, RRMS 1 patients experienced a stable worsening over time, with only 22.2% of worsening patients after 10 years since disease onset.

However, since the definitions of BMS and worsening focus on EDSS and disease duration, the above results do not consider the radiological structure captured by our clustering approach and are thus limited to a clinical differentiation between the two RRMS trends. To also identify physiological differences between the two RRMS trends, we make use of the annualized relapse rate (ARR) of each patient. The main benefit of using the ARR is its ability to both clinically and physiologically quantify the relapsing condition of an RRMS patient, as relapses are defined as periods characterized by clinical deterioration and radiological activity [49]. Thus, the ARR is a way to complement our BMS/non-BMS differentiation from a physiological standpoint. Because of the oscillating trajectories depicted in the non-BMS cluster (Fig. 4b), we expect the non-BMS cluster to be associated with a higher ARR. We observed a mean ARR of 0.26 ± 0.16 within the dark blue cluster and a mean ARR of 0.37 ± 0.27 within the light blue cluster. Additionally, we performed a one-sided MannWhitney U test [50] (*α* = 0.05) to determine whether the ARR distribution underlying the BMS cluster is stochastically less than the ARR distribution underlying the non-BMS cluster (Fig. 4e). The test yielded a *p*-value of 0.01, showing evidence of a difference in ARR between the two clusters. Likewise, Fig. 4f reveals that the non-benign RRMS cluster experiences a significantly higher rate of treatment discontinuation due to the persisting presence of relapses and MS symptoms.

One potential explanation for the existence of the two RRMS sub-clusters is that a substantial portion of the second RRMS cluster could have been misclassified as RRMS instead of SPMS. First, the potential for therapeutic mislabeling still exists, given that the process of verifying it would involve conducting interviews with all the neurologists responsible for diagnosing the patients, which can be a laborious task. Moreover, as of 2023, we have observed three patients from the RRMS 2 cluster transitioning into SPMS, and one patient has passed away. Therefore, it is possible that a few patients from the light blue cluster should have been categorized as SPMS. However, we believe that most of them are more appropriately associated with a distinct sub-group of RRMS, primarily due to important differences in terms of clinical worsening rate (Fig. 4d) and some functional scores such as Ambulation (Fig. 4c, center) which are typically crucial factors in the transition to SPMS [51]. Physiologically, Fig. 4e also shows that the SPMS cluster has a lower ARR, indicating the ongoing presence of a relapsing condition within the RRMS 2 cluster. Thus, our findings support the existence of benign and non-benign forms of RRMS.

To validate these findings, we reproduced our experiments on an independent clinical database collected at the Neuro Rive-Sud clinic in Montreal (see Methods for the details). Overall, the qualitative and quantitative validation results depicted in Fig. S7 and S8 exhibit similar patterns to those obtained with the original data. This gives us confidence that our previous conclusions are general and not an artifact of our data or method.

RF-PHATE has been an essential component of this investigation, and no other methods would have led us to explore these insights. Thanks to its ability to clearly distinguish between the RRMS and SPMS groups (Fig. S4), RF-PHATE enabled us to uncover the high variability within the RRMS cluster of trajectories, prompting us to apply a clustering algorithm to this specific cluster. Additionally, its capability to preserve the structure allowed us to swiftly identify similarities between the lower branch of the RRMS cluster and the SPMS cluster due to their spatial proximity.

### Visualizing A549 antioxidant cell protection via Raman spectroscopy and RF-PHATE

Raman spectroscopy is a non-invasive technology used in various applications of cellular and other biological studies. Raman spectral readings are capable of detecting minute molecular variations and can be used to determine the health of infected or exposed cells [52–55]. These spectral readings often contain noise. Using RF-PHATE and relying on its ability to denoise data as a part of its embedding process, we visually validated the protective and restorative effects of the antioxidants cannabidiol (CBD) and resveratrol (RES) on the health of A549 [55] cells at various levels of exposure to diesel exhaust particles (DEPs), where the supervised labels are the exposure levels. The RF-PHATE embedding shows that the Raman spectra of cells express lower levels of change in the presence of antioxidants when compared to a control group (Fig. 5a,d). Protected cells (Fig. 5b,c,e,f) provide readings more similar to each other but different from the control group (DEP 0) no matter the level of DEP exposure (Fig. 5b,c). This suggests that the antioxidants provide some level of protection against DEP exposure but do not provide total protection. In contrast, PHATE is unable to detect this change as the cells with different exposure levels are difficult to distinguish with or without the antioxidants.

**Fig. 5:**
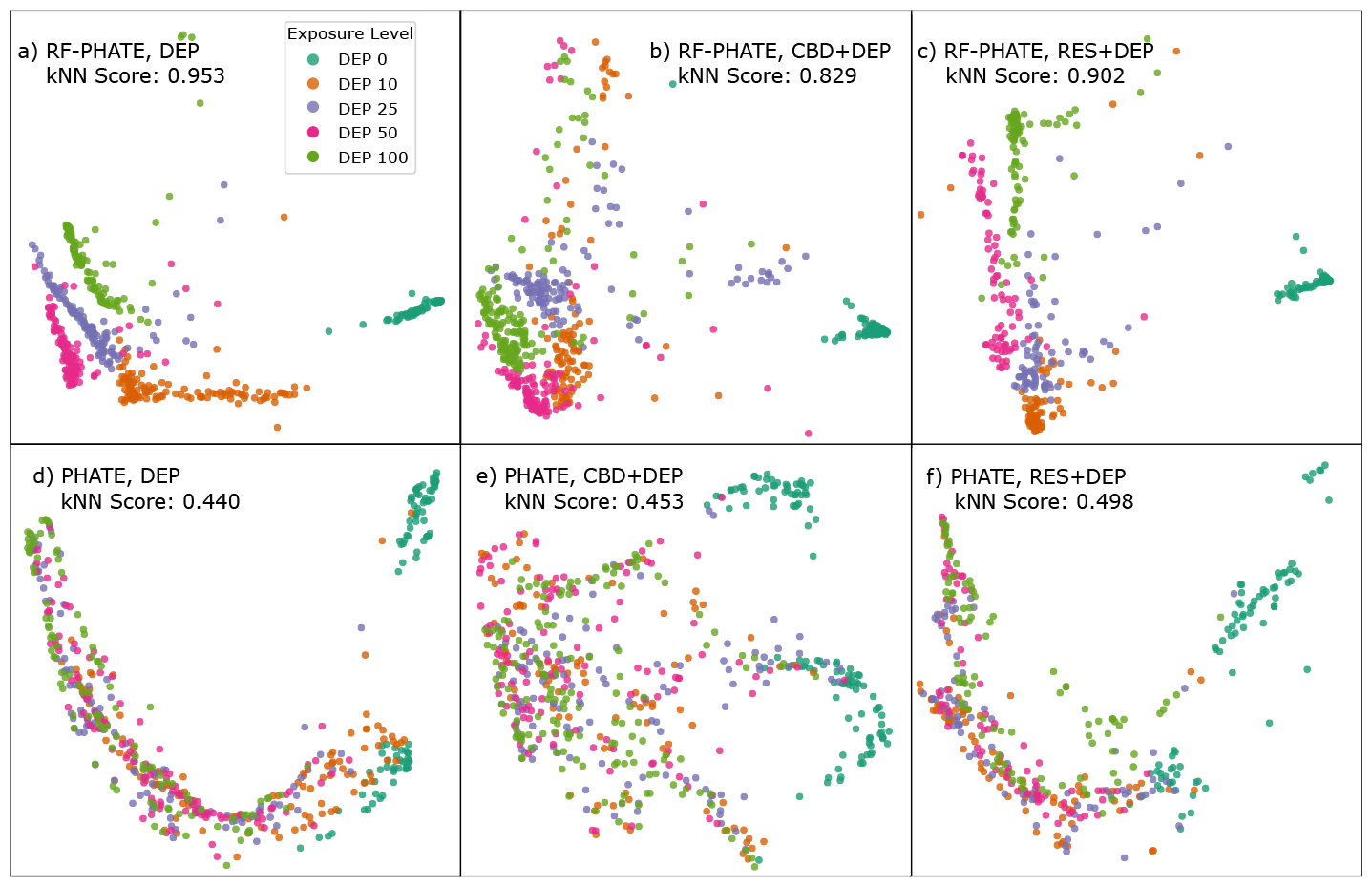
Two-dimensional PHATE and RF-PHATE representations of the collected Raman spectra of A549 cells. Given the DEP exposure values as labels, RF-PHATE shows that the DEP exposure forms partially separable groups. Notably, when no antioxidants (RES or CBD) are present, the levels of DEP exposure are more distinguishable. When antioxidant protectants are added, the control group (no DEP) is distinguishable from the other groups, but exposed groups become less distinguishable from each other, suggesting some level of protection is given at all DEP levels. In contrast, the class separability does not significantly change in the PHATE embeddings (**d, e**, and **f**).

We numerically demonstrate this phenomenon by recording the *k*-NN scores of PHATE and RF-PHATE on the three spectral cell groups exposed to diesel exhaust particles. In each case, the *k*-NN 10-fold cross-validated accuracy means and standard deviations were recorded for the two-dimensional embeddings. The results can be found embedded in Fig. 5. When the cells are treated with CBD, or RES, the accuracy decreases when trained using the RF-PHATE embeddings, numerically showing that cells at all nonzero exposure levels are more similar to each other after treatment. In contrast, the PHATE embedding does not display this pattern.

### Hierarchical visualization of outcomes and immunovirological endotypes in COVID-19 patients

Coronavirus disease 2019 (COVID-19) induces a wide range of patient responses, spanning from asymptomatic cases to severe and fatal outcomes. Plasma viral RNA (vRNA) was quickly identified as a good predictor of COVID-19 patient fatality [56], but the range of patient responses to the disease prompted further investigations into additional immunological features to better understand disease progression. We identified [57] 4 patient endotypes using *k*-means on a set of cross-sectional measurements consisting of plasma vRNA, Spike Receptor Binding Domain (RBD)-specific antibody responses, and plasma levels of cytokines and tissue damage markers. Clusters were visualized using the PHATE embedding of the plasma samples (Fig. 6, a), revealing a global structure primarily driven by antibody and cytokine concentrations [57]. Statistical analyses further confirmed the association between *k*-means clusters and 3 clinically defined patient groups: non-critical survivors (clusters 3 and 4), mechanically ventilated survivors (i.e., critical survivors, cluster 2), and deceased patients (cluster 1).

**Fig. 6:**
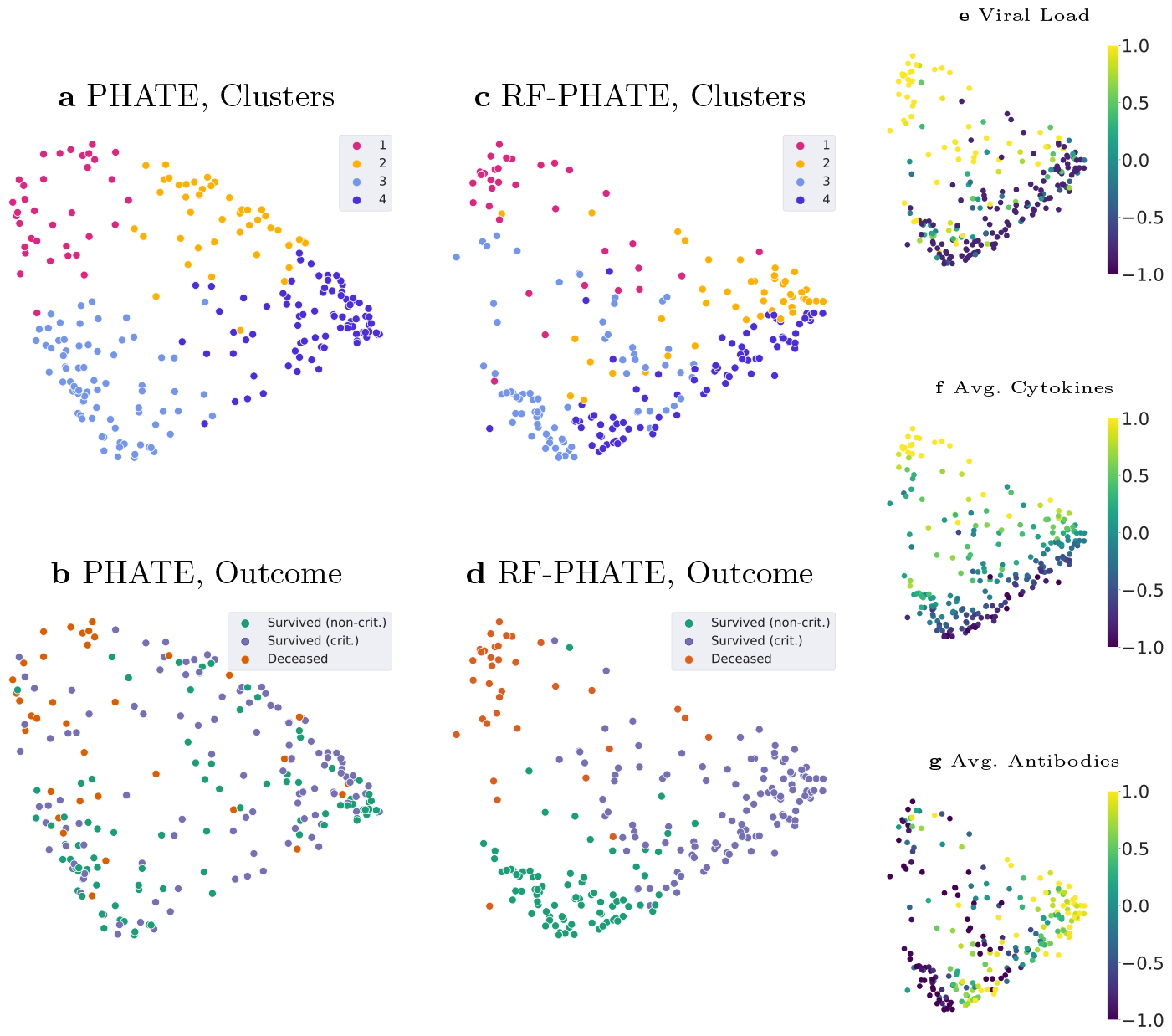
PHATE and RF-PHATE embeddings of plasma samples from hospitalized COVID-19 patients at 11 days after symptom onset (DSO11) [57, 58]. (**a**), PHATE embedding and patient k-means clusters from [57]. The 4 clusters display distinct early immunological profiles and cluster 1 is particularly enriched in high-fatality cases. (**b**), PHATE embedding colored by outcome: survivors are divided into noncritical and critical based on the level of respiratory support they required. (**c**), RF-PHATE embedding using the 3-class outcome as the classification target. The original global geometry of the unsupervised PHATE embedding is still present and some clusters are preserved, illustrating how RF-PHATE factors in both the input data geometry and the supervised signal. (**d**), RF-PHATE embedding colored by the target outcome. As opposed to PHATE, RF-PHATE clearly separates the different outcomes, with both survivor groups located together at the bottom of the embedding. (**e, f, and g**), Average of different input variable categories as coloring for the RF-PHATE embedding. Input variables are standardized (mean 0, variance 1) and the color map is clipped to the range [*−*1, 1] to enhance contrast. The embedding globally correlates with viral load and cytokine concentrations. The heterogeneity of the antibody response in the non-critical survivor group discussed in [57] can be visualized by observing the divide in the lower region of (**g**).

While the associations between clusters and patient outcomes can be measured numerically — e.g., cluster 1 (Fig. 6, a) is particularly fatal —, noise and class imbalance prevent proper visualization of the outcomes (Fig. 6, b). In contrast, RF-PHATE, supervised with the 3-class patient outcome, retains much of the structure present in the original unsupervised setting. E.g., the *k*-means clusters (Fig. 6, c) are still largely separable and are continuously altered to align with patient outcomes (Fig. 6, d), with both survivor groups being located together at the bottom of the embedding. The clean separation between outcomes and the preservation of the original cluster regions demonstrate how RF-PHATE balances both the input data geometry and the guiding target. The overall RF-PHATE geometry separates patient outcomes and globally aligns with vRNA (Fig. 6, e) and average cytokine concentrations (Fig. 6, f). The embedding then exhibits a hierarchical structure: the 2 patient classes are locally organized according to the average antibody concentrations (Fig. 6, g), with a particular “gap” in the non-critical survivor cluster reflecting the heterogeneous antibody profiles of this group as discussed in [57].

Unsupervised embeddings of high dimensional data can identify biologically relevant clusters, such as types of immune response to acute SARS-CoV-2 infection. However, some stratification may be irrelevant in the setting of patient treatment - for example, the distinction between clusters 3 and 4. Supervised embeddings using RF-PHATE have the added benefit of identifying clinically relevant clusters based on an outcome deemed relevant while preserving the structure of the biological data. This type of stratification can be used for early identification of high-risk patients, allowing for treatment prioritization.

We also performed a numerical comparison on patient time series extracted from the COVID-19 data, shown in Table 3. As before, RF-PHATE outperforms all other methods on average across our three metrics.

**Table 3:** Performance scores of 24 different 2D embedding methods on COVID-19 longitudinal data using our three benchmark metrics LGS, LSP, and GSP. The scores are calculated by averaging the results from 20 repetitions. The mean and standard deviations are recorded. Methods are sorted by their average rank across the three metrics (last column). Supervised embedding methods are marked by an asterisk. Just like in our MS comparisons, RF-PHATE is still the best method overall.

| Method | LGS | LSP | GSP | Avg. Rank |
| --- | --- | --- | --- | --- |
| RF-PHATE* | **0.794 ± 0.01** | 0.656 ± 0.01 | **0.745 ± 0.01** | 2.00 |
| ES-ISOMAP* | 0.715 ± 0.00 | **0.757 ± 0.00** | **0.704 ± 0.00** | 2.33 |
| SNMF* | **0.999 ± 0.00** | 0.631 ± 0.00 | 0.651 ± 0.02 | 2.67 |
| SUMAP* | 0.751 ± 0.06 | **0.691 ± 0.11** | 0.575 ± 0.05 | 3.67 |
| SPCA* | 0.313 ± 0.00 | 0.525 ± 0.00 | 0.593 ± 0.00 | 6.00 |
| RF-UMAP* | 0.294 ± 0.01 | 0.510 ± 0.09 | 0.586 ± 0.03 | 7.33 |
| RF-TSNE* | 0.283 ± 0.02 | 0.529 ± 0.06 | 0.557 ± 0.05 | 8.00 |
| PLS-DA* | 0.312 ± 0.00 | 0.457 ± 0.00 | 0.486 ± 0.00 | 8.67 |
| RF-ISOMAP* | 0.334 ± 0.02 | 0.349 ± 0.06 | 0.506 ± 0.11 | 9.33 |
| PHATE | 0.270 ± 0.00 | 0.596 ± 0.01 | 0.436 ± 0.00 | 9.33 |
| S-TSNE* | 0.600 ± 0.01 | 0.426 ± 0.03 | 0.135 ± 0.04 | 10.33 |
| ISOMAP | 0.276 ± 0.00 | 0.389 ± 0.00 | 0.421 ± 0.00 | 12.00 |
| UMAP | 0.233 ± 0.01 | 0.411 ± 0.08 | 0.255 ± 0.11 | 14.00 |
| NCA* | 0.286 ± 0.00 | -0.150 ± 0.05 | -0.179 ± 0.03 | 15.33 |
| RF-MDS* | 0.196 ± 0.00 | 0.289 ± 0.06 | 0.311 ± 0.03 | 15.33 |
| LLE | -0.094 ± 0.00 | 0.439 ± 0.00 | -0.336 ± 0.00 | 17.67 |
| ESLLE* | -0.431 ± 0.01 | 0.250 ± 0.11 | 0.245 ± 0.02 | 17.67 |
| PCA | 0.257 ± 0.00 | -0.296 ± 0.00 | -0.214 ± 0.00 | 18.00 |
| T-SNE | 0.219 ± 0.00 | -0.158 ± 0.02 | -0.175 ± 0.01 | 18.00 |
| MDS | 0.238 ± 0.00 | -0.327 ± 0.01 | -0.197 ± 0.03 | 18.33 |
| DM | -0.726 ± 0.00 | 0.136 ± 0.00 | -0.150 ± 0.00 | 19.00 |
| LAP. EIG. | 0.234 ± 0.00 | -0.539 ± 0.00 | -0.382 ± 0.00 | 20.33 |
| KPCA | 0.186 ± 0.00 | -0.511 ± 0.00 | -0.389 ± 0.00 | 21.67 |
| KSPCA* | 0.175 ± 0.00 | -0.814 ± 0.00 | -0.729 ± 0.00 | 23.00 |

## Discussion

When working with high-dimensional biological or other complex data, a lower-dimensional visualization of the data can assist practitioners in better understanding the relationships between measured variables and a labeled response. However, the majority of existing DR approaches do not use extra, expert-provided auxiliary information to guide the embedding generation process. Previous existing supervised methods distort the global data structure by imposing class-driven constraints that artificially separate observations of different classes in the low-dimensional embedding. In this paper, we provided RF-PHATE as an alternative to these unsatisfactory supervised methods. RF-PHATE combines the powerful predictive power of random forests with diffusion-based informational geometry designed specifically for visualization. RF-PHATE is scaleable to small and large datasets, employs the flexibility of random forests, and works well for both continuous and categorical labels. RF-PHATE is robust to noise due to both the ability of random forests to determine feature importance and because of its diffusion dynamics.

We showed the ability of RF-PHATE to produce interpretable insights in a longitudinal study of MS patients from which we uncovered a non-benign sub-group of RRMS patients. RF-PHATE demonstrated superior performance compared to existing DR methods in embedding MS follow-up data by effectively separating the two RRMS and SPMS groups while preserving the essential structure of the original space. In addition, RF-PHATE proved useful in practice by accomplishing two objectives: retrieving domain knowledge of RRMS and SPMS courses based on clinical and radiological features, and highlighting a non-benign RRMS sub-group that was not explicitly identified in our initial database. The non-benign RRMS sub-group does not primarily result from SPMS patients being misclassified. Instead, compared to SPMS, this RRMS sub-group exhibits a lower worsening rate (or “silent progression”) and a higher ARR showing an active relapsing-remitting condition. This conclusion strengthens the notion of a distinct RRMS sub-group with unique characteristics and specific needs.

Given the significant treatment failure observed in the non-benign RRMS cluster (Fig.4f) despite the use of similar medication (Fig.S6), our future focus involves prioritizing attention on RRMS patients whose disease course aligns with non-benign RRMS cluster. This shift aims to facilitate early and tailored treatment for improved patient outcomes. Additionally, the inclusion of spinal cord MRI data may enhance the visual distinction between MS subgroups, considering the substantial impact of spinal cord lesions on clinical symptoms [59, 60]. Furthermore, we may explore the reclassification of RRMS patients into SPMS or other RRMS sub-types to enhance the efficiency of downstream prediction tasks related to the progression of MS. By specifically targeting three distinct MS profiles or degrees of severity—RRMS 1, RRMS 2, and SPMS—this approach enables an early prognosis of the forthcoming MS course.

We demonstrated RF-PHATE’s ability to create meaningful visualizations in a variety of biological contexts, providing new insights or corroborating domain expert knowledge. In the noisy context of Raman spectroscopy, RFPHATE displayed the protective effects of two types of antioxidants against diesel exhaust particles on A549 cells. In a COVID-19 case study, we showed that RF-PHATE embeddings aligned known geometric data structures with target labels to produce a visualization that both corroborates previous findings of covariates associated with patient outcomes and enriches these relationships in a hierarchical structure with increased interpretability. We anticipate that RF-PHATE is capable of producing valuable insights in a variety of fields and applications, both biological and otherwise. We expect to see new uses in private research and industry, with biomedical and clinical data, by revealing information in these fields via human-interpretable visualizations.

## Methods

Here we provide additional details concerning our methods including an overview of supervised manifold learning, background information on random forests, the RF-GAP proximities in manifold learning, descriptions and benefits of diffusion-based information geometry, further comparisons on data with noisy variables, details on our measures for quantifying embedding fit, further historical context for the EDSS score, and specific methodologies for data preparation and analysis including the use of dynamic time warping to adapt RF-PHATE to time series data.

### Overview of supervised manifold learning

Visualizing high-dimensional data often serves as an early step in exploratory data analysis. Complex relationships between data points and features become more meaningful when practitioners can view relevant spatial or temporal representations of the data as a complement to summary statistics. Manifold learning provides a means to generate low-dimensional embeddings that can serve as tools for high-dimensional data visualization or data preprocessing.

Most manifold learning approaches do not use auxiliary information, such as labels, in the embedding-learning task. It is reasonable to ask if/how labels can be used in this traditionally unsupervised machine-learning task. Expert-provided labels should provide additional insights into the data structure relative to the label information. Providing an algorithm a target to seek (i.e., supervised learning) can assist the algorithm in learning a representation from features that relate to the supervised task.

Supervised counterparts for common manifold learning algorithms exist and are typically adapted at the algorithm’s distance- or affinity-learning step. In some cases, distances are rescaled depending on class affiliation, as in the case of WeightedIso [61], a supervised version of Isomap [5]. Others additively increment distances between points of opposing classes, as is done in SLLE [11], a supervised version of local linear embedding (LLE) [4]. In the cases of supervised (S)-Isomap [62], enhanced supervised (ES)-Isomap [9], enhanced supervised (ES)-LLE [10], supervised Laplacian Eigenmaps (S-LapEig) [63], and supervised (S)-tSNE [8], dissimilarities are derived as class-conditional adaptations of a Gaussian kernel function.

However, manifold learning methods using these class-conditional dissimilarities suffer from a number of weaknesses: (1) The global structure relative to inter-class relations is disrupted. This is corroborated by the authors of [12, 64], who show these dissimilarity functions are order-non-preserving isomorphisms and should not be used for visualization tasks lest one draws misleading conclusions about the data structure. (2) The utility of downstream tasks is diminished. For example, class-conditional dissimilarities often induce perfect class separation which is inconsistent with the intrinsic dataset structure. Classification following DR in this manner can give unrealistically low error rates. (3) These dissimilarity measures do not provide a natural extension for continuous-valued labels and are thus incompatible with regression problems. (4) Dissimilarities between labeled data and unlabeled data (e.g., a training set vs. a test set) are not defined.

To counter each of these weaknesses, we designed RF-PHATE to use the random forest-based RF-GAP proximities [14] as a similarity-learning step instead of conditionally increasing distances between observations of different classes. RF-GAP similarities are computed in a manner that preserves random forest learning and thus provide a powerful local similarity representation for supervised manifold learning.

RF-GAP proximities serve as similarity measures that use data labels in accordance with the supervised model’s learning and thus comprise a measure that overcomes the weaknesses of the class-conditional distances: (1) RF-GAP proximities form a local similarity measure that partially retains global information through the forest’s recursive splitting process. Thus, the data geometry learned by the random forest captures hierarchical feature importance and does not induce artificial class separation but distinguishes between classes in a manner consistent with the intrinsic structure of the dataset relative to the labels. Thus instead of distorting the global structure, the proximities naturally retain observational relationships. (2) Rather than diminishing the integrity of downstream tasks, RF-GAP proximities retain random forest learning without overfitting to the training data. Thus, downstream tasks are positively influenced by this supervised information. (3) Class-conditional similarities are subject to discrete labels, but random forests can handle both discrete and continuous labels. RF-GAP proximities provide a way to perform supervised manifold learning with continuous labels. (4) Class-conditional similarities are not extendable to an unlabeled test set, but a trained random forest can define similarities to out-of-sample observations, supporting semi-supervised learning and other tasks.

### Similarity learning with random forests

Though dozens of variations have been introduced since the random forest’s conception in 2001, the original random forest algorithm [13] is still used in a variety of applications today. Citing a few recent examples, random forests have been used to classify water salinity [65], groundwater nitrate pollution [66], to assess the risk of coronary heart disease [67], to predict cellular types and health using Raman Spectroscopic features [55], to classify oncosomatic gene variants using next-generation sequencing, outperforming recurrent neural networks [68], to predict compounds with anti-aging properties [69], and to predict the prognosis of diabetic patients [70].

In addition to prediction problems, random forests are also commonly used for feature selection or data preprocessing. Feature analysis for traditional ecological models was performed using random forests in [71]. Longitudinal predictors of health in exposome studies were identified using random forests in [72]. Feature selection has also been done via random forests in the context of surface-enhanced Raman scattering data [73], genome-wide associations using population structures and individual genetic variants [74], and for deep-learning preprocessing to prevent overfitting in gene-expression classification [75].

Random forest usage is universal because they are simple to use and they provide a broad array of benefits in addition to prediction. In particular, random forests perform well with little or no parameter tuning and can be used when the number of features exceeds the number of observations (*p > n*), which is often the case with biological data. They are compatible with both regression and classification problems and can be applied to both continuous and categorical features without feature engineering. This last benefit allows forest-based similarities to construct meaningful relationships with observations of mixed feature types which is often a factor in clinical studies [76]. In contrast, most manifold learning methods do not naturally handle mixed features.

In simple terms, random forests are an ensemble of randomized decision trees. Each decision tree is formed using a random bootstrap sample (sampled with replacement) of training data. In the literature, observations kept in the bootstrap sample are called in-bag observations, while those not included are called out-of-bag (OOB) observations. The OOB observations serve as a quasi-test set for model validation.

The trees are grown by a recursive splitting of the data across a randomized selection of features in a manner that maximizes purity within resulting nodes. The recursive process continues until a stopping criterion is met. The set of terminal nodes forms the decision space of the random forest. A new observation, *x*_0_, is passed down each decision tree and a vote takes place within its terminal node of residence. An overall, unweighted vote is counted across all trees in the forest, providing the predicted label for *x*_0_. It is important to note that the collection of in-bag observations in the terminal nodes of *x*_0_ exclusively determines its predicted label. Thus, the random forest can be viewed as a nearest-neighbor decision algorithm, where “neighbors” are in-bag observations in a shared terminal node [77].

The space of terminal nodes or “neighborhoods” can be used to define a similarity or proximity between observations. Since observations are organized according to splits across the features that are most useful for partitioning data, these similarities naturally incorporate feature importance. While two observations may be considered distant in a Euclidean space, they might be very similar with regard to important features for the supervised problem. Random forests are also robust to noise, making forest-derived proximities useful to process noisy, high-dimensional data.

Random forest proximities are typically constructed pair-wise according to terminal node residence. Random forest-based proximities were first defined as the proportion of trees in which a pair of observations shares a terminal node [13, 78]. However, this proximity definition tends to exaggerate class distinction as in-bag examples of opposing classes necessarily reside in different terminal nodes when trees are fully grown. As a fix, proximities are often constructed using only OOB pairs of points [79, 80]. These OOB proximities are still insufficient at capturing the random forest’s learning because 1) they do not directly use training examples (in-bag observations), and 2) they fail to account for the number of observations in each terminal by which random forest voting is conducted [14].

To fully incorporate the random forest’s learning and yet prevent overfitting to the training data, the proximities need to be constructed in a manner consistent with the random forest’s modus operandi: predictions for new or OOB observations are made by majority vote or by averaging labels of in-bag observations within each terminal node. That is, instead of counting pairwise interactions without regard to other points in a common terminal node, the proximities should be formed relative to the number of in-bag observations within the terminal nodes. Rather than constructing proximities between training points, the training points are used to construct proximities to OOB points. In this way, the proximities to a given OOB observation are consistent with the forest’s predictive model. It was shown in [14] that thus defined RFGAP proximities are capable of perfectly recreating the random forest OOB and test prediction when serving as weights in a weighted average (or weighted majority vote) of the corresponding data labels. It was also shown that multiple proximity-based applications are improved by using RF-GAP proximities. Thus, we use them as a supervised similarity measure for supervised manifold learning.

A few modifications of the RF-GAP proximities are required to use them in manifold learning. First, by default, the diagonal entries of the RF-GAP proximities are defined to be zero as an observation cannot be simultaneously in-bag and out-of-bag for a given decision tree. To overcome this, we redefine the diagonal entries by, in essence, passing an identical, OOB point down each tree in which the original point is in-bag. Using this point to generate the diagonal proximities we maximize the similarity between an observation and itself while keeping these diagonal entries on a similar scale to the rest of the proximities. Additionally, RF-GAP proximities were designed to be capable of reconstructing the random forest predictions by using them as weights. Thus, they are not symmetric, although it was demonstrated in [14] that RF-GAP proximities are asymptotically symmetric as the number of trees increases. In applications requiring symmetric affinities, this can easily be fixed by symmetrizing the proximities. With these adjustments, we can use RF-GAP proximities as a supervised kernel to be used in manifold learning.

### Diffusion-based information geometry

Early attempts at diffusion-based dimensionality reduction, such as Diffusion Maps [24], use a *k*-NN graph upon which a kernel function, such as a Gaussian kernel with a fixed bandwidth, is applied to represent observational similarities. The similarity matrix is row-normalized to form a stochastic matrix, *P*, known in this context as the diffusion operator. The diffusion operator represents the probabilities of all possible single-step transitions between observations, where high affinities correspond to high transition probabilities. Through a powering process, *P*^*t*^ represents a random walk across the manifold structure where global relationships are learned. During this process, small probabilities are quickly reduced to zero, thus filtering or denoising the learned manifold. Finally, dimensionality reduction can be applied (e.g., eigendecomposition) to map to lower dimensions.

Although this diffusion process is intuitive and suitable for dimensionality reduction, some weaknesses need to be addressed to create meaningful visualizations. First, a fixed kernel bandwidth for all points is often not appropriate as the data may not be sampled uniformly. Second, choosing a good time scale *t* for the diffusion process is difficult and largely overlooked in classical diffusion maps. A small value of *t* can lead to insufficient denoising and an overemphasis on the local structure while a large value of *t* can lead to oversmoothing and an overemphasis on the global structure. Third, the eigendecomposition in Diffusion Maps often places the information learned in *P*^*t*^ into different and higher dimensions [1, 81], which is not amenable for visualization.

Instead of applying a kernel function to Euclidean distances, we encode local similarities using RF-GAP proximities. These locally-adaptive similarities contain local to long-range dependencies and overcome the weakness of a fixed-bandwidth kernel function. However, if the data has many disjoint clusters or clusters with limited connections, global relationships may not be properly reflected in the transition process. For example, the transition process may favor a single cluster with only transition inlets but no outlets. We counter this scenario with the introduction of a damping factor, *β* ∈ (0, 1], inspired by Google’s PageRank algorithm which is used to rate website importance [15].

The PageRank damping factor was inspired to overcome the problems of “traps” and “dead ends”, that is, sets of websites from which all outward links point towards the same group of websites or towards a single external webpage. Analogously, we counter poor exit probabilities by redefining the diffusion operator *P* as 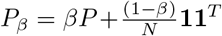, where **1** is a vector of ones of length *N* and *β* is typically set between 0.8 and 0.9. Here, the transition probabilities allow for random jumps creating small exit probabilities from isolated clusters or dead-ends. This has the effect of maintaining global structure in the presence of disjoint clusters.

To properly select *t*, we follow the inspiration of PHATE [1], which uses von Neumann Entropy (VNE) of the diffused operator to provide a good choice of *t* for visualization. The VNE is a soft proxy for the number of significant eigenvalues of 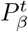. Typically, *t* is chosen to be around the transition from rapid to slow decay in the VNE as this is considered to be a point in the diffusion process where noise has been eliminated and oversmoothing begins [1].

To overcome the weaknesses stemming from the eigendecomposition in standard Diffusion Maps, we apply an information distance [16] to the powered diffusion operator 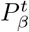 to create a form of potential distance [1]. The potential distance is sensitive to differences in both the tails and the more dense regions of the diffused probabilities, resulting in a distance that preserves both local and global relationships. These distances are then embedded into low dimensions using metric MDS. This embeds the information in low dimensions for better visualization. The full RF-PHATE algorithm is given in Algorithm 1.

#### Algorithm 1

The RF-PHATE algorithm

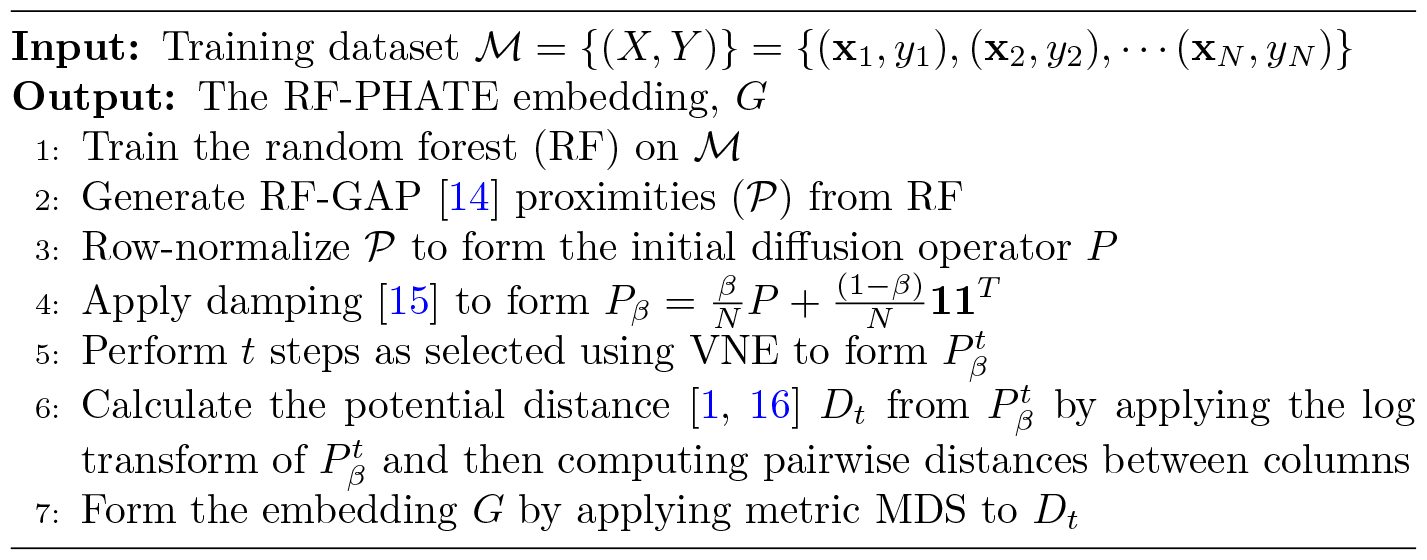

### Embedding noisy data

Expert-generated data labels assist the learner in knowing what types of information to seek, but not how to perform the learning. In some cases, relationships between the inherent data structure and the data labels may not be trivial. In others, excessive noise in the data can prevent the learning of meaningful structures without the assistance of data labels. In Fig. 3, we showed how RF-PHATE is able to ignore irrelevant (i.e., noise) features and capture the true structure of the artificial tree data, while other methods fail in the presence of noisy features.

Here we provide an additional example of a dataset with well-known structures to which we intentionally add noise features. With these examples, we demonstrate the ability of RF-PHATE to uncover a meaningful data structure relative to the labels in the presence of noise. This example presents a noised version of the simple and well-known Iris dataset [82]. In this example, we combined the original dataset (with features rescaled from 0 to 1) with an additional 500 uniformly sampled 0-1 noise features. The results are shown in Fig. S1. In this scenario, unsupervised approaches are incapable of determining a meaningful embedding. Each of the compared unsupervised methods generated a seemingly random cloud of points. Class-conditional supervised methods all tend to completely separate points of different classes, which is contrary to the ground-truth structure, in which there is a small amount of overlap between the Versicolor and Virginia species. In contrast, RF-PHATE preserves this structure even in the presence of noisy features.

### Quantifying supervised embedding fit

Traditional tools for assessing DR quality are insufficient in a supervised setting and are not necessarily intended to provide a quantitative assessment of the embedding’s useability for visualization. For example, MDS [17] seeks to minimize stress, the normalized squared differences between distances in the original space and the embedding space, but low stress does not necessarily correlate with the visual quality of the embedding [83]. To quantify the quality of supervised embeddings, one approach is to record the accuracy of a *k*-NN predictor trained on the embedding. However, this favors methods that emphasize class separation rather than structure preservation. Furthermore, perfect class separation can be attained by supervised manifold learning approaches using class-conditional distances (Fig. S1 and Fig. 3, for example).

**Fig. S1:**
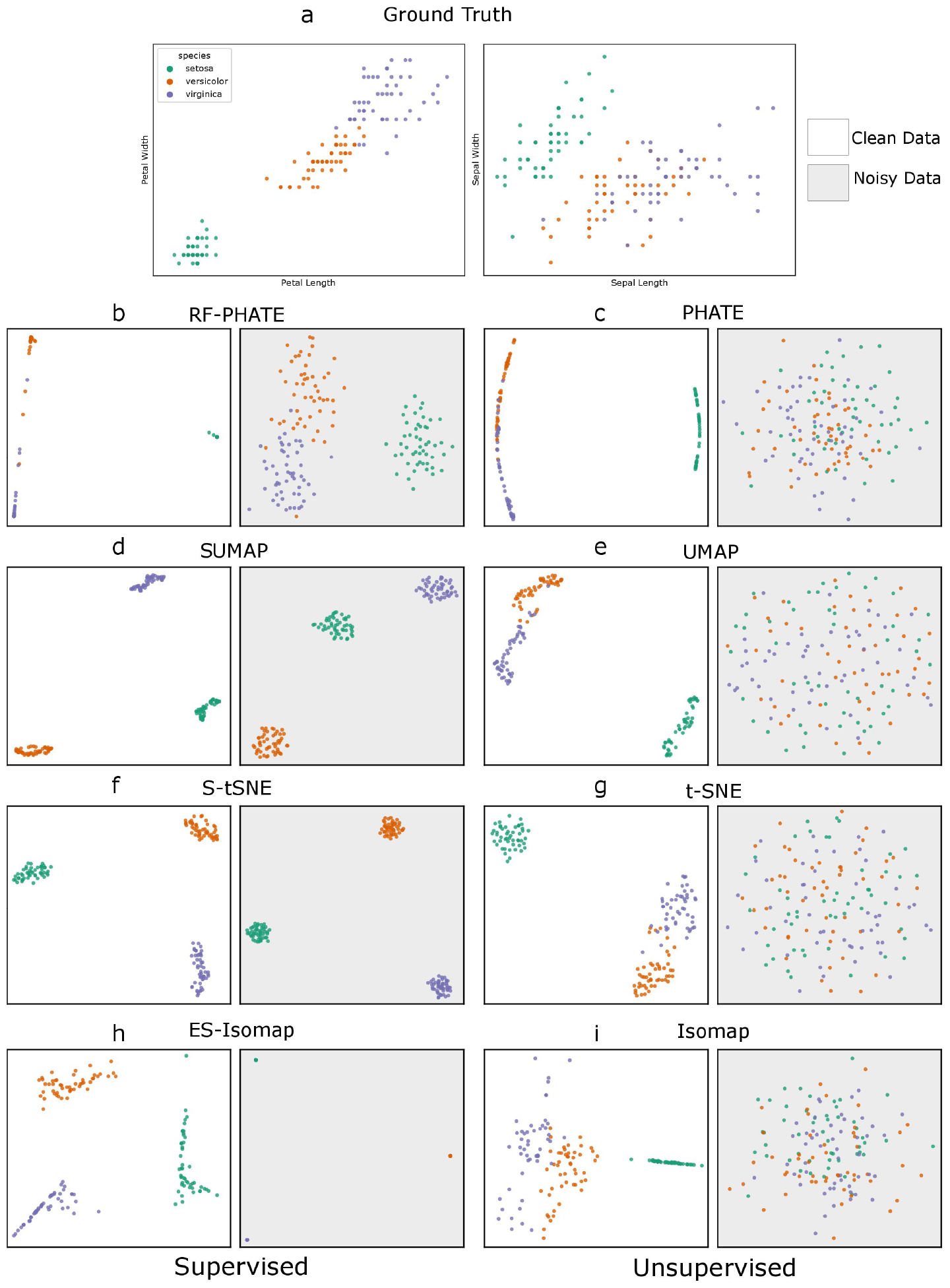
Here we added 500 noise variables (each uniformly distributed from 0 to 1) to the well-known Iris dataset [82]. **a**, presents two of the pairwise variable plots, each of which demonstrates the separability of the Setosa class from the other two, while simultaneously showing that the other two classes have some overlap. **c, e, g, i**, unsupervised methods can generally reconstruct a meaningful representation in the absence of noise (white background) but are incapable of showing these relationships with noise (gray background). **d, f, h**, class-conditional supervised methods tend to completely separate the classes, which is not consistent with the known data structure. On the other hand, RF-PHATE, **b** preserves the inherent structure despite the noise.

To be useful in exploratory analysis, the goal of our method is not explicitly to separate classes but to uncover the intrinsic data structure relative to the supervised task. We thus use three different measures to quantify an embedding’s goodness of fit. Low-dimensional Group Separation (LGS) assesses the extent to which the embedding is consistent with the classification problem; this method uses the silhouette score of the embedded points using data labels in lieu of its traditional use with cluster labels. The Local Structure Preservation score (LSP) correlates the local feature importance of the supervised task with the feature importance for the embedding construction. This measures the extent to which the embedding can be recovered using the data features pertinent to the classification or regression problem at a local level. Global Structure Preservation (GSP) correlates the pairwise distances of each univariate feature with the pairwise distances of the embedding. The degree to which these correlations relate to the global feature importance of the original prediction problem determines the extent to which the global structure is maintained in low dimensions relative to the prediction problem. Further details on these metrics are provided below:

a. **LGS**: the **L**ow-dimensional **G**roup **S**eparation score is defined as the overall silhouette score of the grouping in the embedding space determined by the label assignment **Y**:

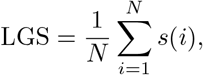

where
  − 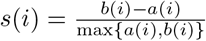 is the silhouette score of the *i* embedded point,
  − *a*(*i*) is the average distance between embedded point *i* and all other embedded points belonging to the same class,
  − *b*(*i*) is the average distance between embedded point *i* and all other embedded points in the other class.
  The overall silhouette score measures clustering quality [84] which is commonly used in the exploratory analysis of healthcare data [85–87]. It also has the convenience of ranging between − 1 and 1, where values near 1 indicate that objects are very well matched to their own group and poorly matched to neighboring groups. Values near 0 indicate that objects are equally well matched to their own group and to neighboring groups, and values near − 1 indicate that objects are poorly matched to their own group and well matched to neighboring clusters. Instead of evaluating a clustering algorithm, we use the overall silhouette score to evaluate a DR method in separating the existing groups in the embedding space to provide the user with a visualization that clearly highlights class relationships. LGS does not have any tunable parameters.
b. **LSP**: the **L**ocal **S**tructure **P**reservation score [88] is used to determine how important features are used in the embedding construction. In a supervised context, features important for classification or regression should dominate the low-dimensional local structure of the embedding. Thus, we produce two rankings of the input features with respect to (1) their importance to the classification or regression task and (2) their importance to the local embedding structure generation. A good embedding should preserve the respective feature rankings. LSP uses a trained prediction model (typically *k*-NN) to assess feature importance scores using randomized permutation importance, *𝒞*, for the original prediction task. A similar model is trained to regress onto the embedding and feature importance scores, *ℒ*, for this task are also assessed. LSP is calculated as the correlation between these two sets of feature importance scores, that is,

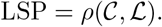

This correlation score also ranges from − 1 to 1, like LGS, allowing us to use them as a benchmark for a direct comparison between embedding algorithms.
c. **GSP**: **G**lobal **S**tructure **P**reservation score is computed similarly to LSP. A model’s feature importance scores, *𝒞*, is correlated with another set of feature importances *𝒢* related to the global embedding structure generation.

Becht et al. [89] demonstrated that UMAP preserves global structure better than *t*-SNE by correlating pairwise distances in the original space with pairwise distances in the embedding space, where a high correlation is assumed to indicate good preservation of the original data structure from a macroscopic point-of-view. Here, we modify their approach to incorporate our hierarchical structure preservation with respect to feature importance for the prediction task. Instead of directly computing pairwise distances in the original, high-dimensional space, we compute pairwise distances within each univariate feature space and correlate each of the corresponding condensed pairwise distances 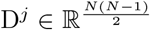 with the condensed pairwise distances 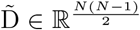 between the embedded space 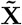. Thus, the quantity 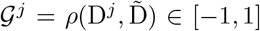 measures the importance of feature *j* in the global structure generation, and the set of global feature importances is defined as *𝒢*= {*𝒢*^*j*^ : *j* = 1, …, *K*}. The GSP score of a DR method is then defined as the following Spearman’s rank correlation coefficient:

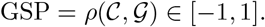

We adapt each of these scores to time series data as described later in the Methods.

### EDSS historical context

Accurately classifying the course of MS into sub-types is very important for communication, prognostication, design and recruitment of clinical trials, and treatment decision-making [90]. To this end, Lublin et al. [90] proposed a baseline classification of MS into four major forms: Clinically Isolated Syndrome (CIS), Relapsing-Remitting MS (RRMS), Secondary-Progressive MS (SPMS) and Primary-Progressive MS (PPMS). Two main descriptors underlie these phenotypes: disease activity, as evidenced by relapses and/or new activity on magnetic resonance imaging (MRI), and progression of disability.

However, retrospective evaluations of physicians are inherently subjective and limited in terms of processing information. Additionally, the course of MS is widely recognized as highly heterogeneous and difficult to predict on an individual basis over time [91]. As a result, historical values of the expanded disability status scale (EDSS), computed as a nonlinear combination of eight functional scores [25, 92], remain the gold standard to characterize MS progression and relapses [26].

While other available historical data, such as the eight underlying functional scores or imaging features, are often not explicitly used by physicians in the MS diagnosis, our study is motivated by the assumption that there is an implicit interest in integrating these features to unveil more specific sub-types of MS. For instance, according to Lublin et al. [90], MRI data can help in determining active and inactive sub-types of RRMS and SPMS through contrast-enhancing lesions or new or unequivocally enlarging T2 lesions assessed at least annually with the aim of potentially switching to a high efficacy disease-modifying therapy and reduce the risk of relapse in case of high disease activity [93]. This also led researchers to classify MS patients using features that better reflect the pathophysiology of the disease, such as MRI lesions [94], underlying biological mechanisms, or the continuous aspect of disease progression [95]. Still, there is a lack of consensus regarding the clinical significance of MRI data as well as the clinically important changes that may result from changes in MRI data [96]. This means that, while additional features may be useful, EDSS remains the main differentiating factor between MS sub-types at this point.

### MS data collection and description

The MS data was obtained from patients diagnosed with clinical and neuropathological MS diagnosis according to the revised 2010 McDonald’s criteria [45]. Disability was scored by a qualified neurologist using the EDSS [25]. More specifically, our raw longitudinal database consists of two distinct MS follow-up data sets. The clinical set consists of 16,371 clinical visits with 636 distinct patients from which 9 features (Table S1) quantifying disability in MS were collected from a clinical examination. The radiological set consists of 6,327 radiological visits across 1,504 patients from which 6 radiological features related to the chronic inflammatory nature of the disease were recorded from MRI sessions. For a more detailed description of the available input features and their calculation, refer to Tables S1 and S2 as well as the guidelines of the reference papers [25, 92]. Note that both databases can be expressed as a set of time series from each patient containing the history of their clinical and radiological visits.

**Table S1:**
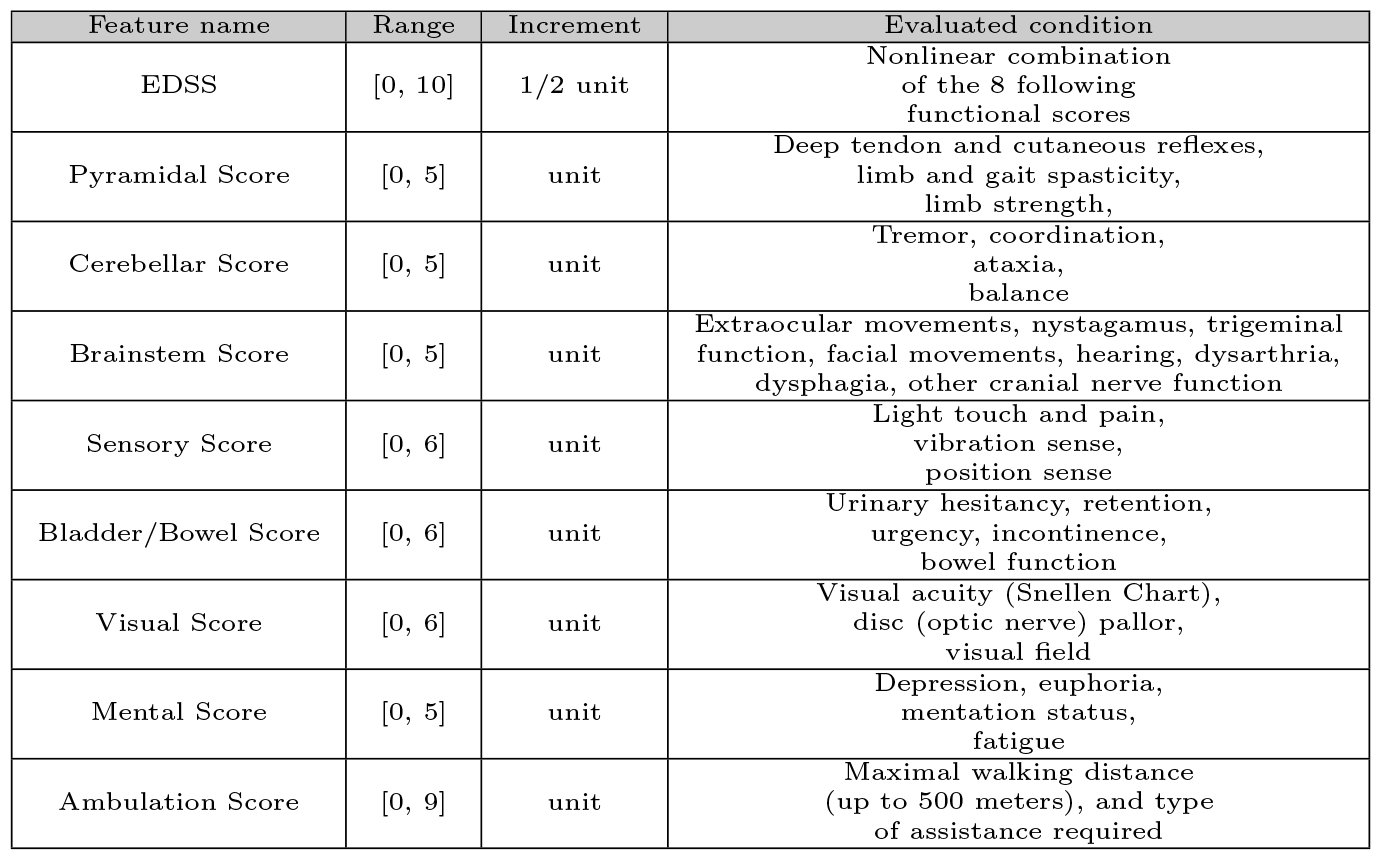
Description of the 9 longitudinal input features recorded for each patient in the clinical database. Higher values indicate worse patient conditions.

| Feature name | Range | Increment | Evaluated condition |
| --- | --- | --- | --- |
| EDSS | [0, 10] | 1/2 unit | Nonlinear combination of the 8 following functional scores |
| Pyramidal Score | [0, 5] | unit | Deep tendon and cutaneous reflexes, limb and gait spasticity, limb strength, |
| Cerebellar Score | [0, 5] | unit | Tremor, coordination, ataxia, balance |
| Brainstem Score | [0, 5] | unit | Extraocular movements, nystagamus, trigeminal function, facial movements, hearing, dysarthria, dysphagia, other cranial nerve function |
| Sensory Score | [0, 6] | unit | Light touch and pain, vibration sense, position sense |
| Bladder/Bowel Score | [0, 6] | unit | Urinary hesitancy, retention, urgency, incontinence, bowel function |
| Visual Score | [0, 6] | unit | Visual acuity (Snellen Chart), disc (optic nerve) pallor, visual field |
| Mental Score | [0, 5] | unit | Depression, euphoria, mentation status, fatigue |
| Ambulation Score | [0, 9] | unit | Maximal walking distance (up to 500 meters), and type of assistance required |

**Table S2:** Description of the 6 longitudinal input features recorded for each patient in the radiological database.

| Feature name | Domain | Description |
| --- | --- | --- |
| Nb Lesions T2 | Any integer | Lesion count using the T2 imaging method [97] |
| Nb New or Enlarging Lesions | Any integer | New/enlarging lesion count with reference to a baseline scan |
| McDonald Lesions T2 | 0, 1–2, 3–8, 9+ | T2 lesion count with categories according to the McDonald criteria |
| McDonald Lesions Infratentorial | 0, 1+ | McDonald lesion count located in the infratentorial brain region |
| McDonald Lesions Juxtacortical | 0, 1+ | McDonald lesion count located in the juxtacortical brain region |
| McDonald Lesions Periventricular | 0, 1–2, 3+ | McDonald lesion count located in the periventricular brain region |

At the end of the follow-up study, each patient was assigned one of the four typical MS types: Clinically Isolated Syndrome (CIS), Primary Progressive MS (PPMS), Relapsing-Remitting MS (RRMS), or Secondary Progressive MS (SPMS). According to the official classification provided by Lublin et al. [90], RRMS is a type of MS with relapses, defined as bouts of clinical deterioration (appearance of new symptoms or worsening) for a period of 24 hours or more—in the absence of an infection or fever. Relapses in RRMS are followed by a variable degree of recovery, without overall disease worsening outside of relapses. This form of MS can (but not always) progress to SPMS during which patients experience insidious accumulation of neurological disability outside of relapses. In that case, the SPMS type is assigned to the patient. Since CIS and PPMS are less common forms of MS [90], we will be focusing on RRMS and SPMS throughout this study.

In addition, the collected data offers contextual information that can be valuable for correlating with our main longitudinal features and RRMS/SPMS classes. The worsening _*i*_ of an MS patient after a period of *𝒲*_*i*_ years is a clinically motivated binary (0/1) worsening score, defined from EDSS as

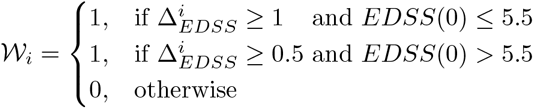

where 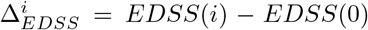 denotes the difference between the EDSS after *i* years and the EDSS at the beginning of the disease course. In the database, the worsening is given for *i* = 1, 2, 5, 10 years. The annualized relapse rate (ARR) is computed for each patient as the total number of relapses divided by the number of years since disease onset. In many clinical trials, it is a primary endpoint, because relapses are meaningful as they reflect how individuals experience fluctuations in symptoms that are central to relapsing MS.

Lastly, our data set contains typical demographic characteristics and some treatment details, including the duration of MS-specific medication usage, along with the reasons for discontinuation. Table S3 presents a summary of some patients’ characteristics within the two RRMS and SPMS classes after data imputation and processing. As expected, the SPMS group exhibits a greater EDSS on average, as well as higher worsening rates. While the rate of relapses generally declines in SPMS patients, the similar ARR observed in Table S3 for both groups primarily results from the fact that SPMS patients inevitably experience RRMS before progressing to SPMS. This means some patients might have transitioned at a later stage or encountered high ARR during their RRMS phase, thereby contributing to an elevated ARR in the long run.

We also include the Spearman correlation matrix of the input features in Fig. S2.

### MS data imputation and processing

We impute missing disability and MRI feature values using linear interpolation. For ordered categorical features, we first associate each category with an integer {0, 1, 2, …} and round the imputed value to the nearest integer. To form a single database describing patients’ follow-up in terms of both disability and MRI features, we linked clinical visits to MRI visits that happened no more than six months apart. Clinical visits without MRI data falling into that time period are simply removed. Afterward, we re-sample the combined dataset on a 6-month frequency and normalize the time between patients by setting the origin to the date of MS disease onset (ddo). In other words, the time (ddo) of a patient’s visit corresponds to the number of days between the date of visit and the date of MS disease onset.

Since none of the DR methods presented in this work directly receive timeseries objects as input, we also consider the normalized time component as an input feature to take the temporal structure of the input data into account, giving a total of *K* = 16 input features for each patient. Finally, we standardize features and remove patients without the available target of interest. That is, we removed the seven patients diagnosed with Clinically Isolated Syndrome (CIS) or Primary Progressive MS (PPMS), which are less common forms of MS. This pre-processing results in *M* = 5,378 visits, including both clinical and radiological features, spread across *N* = 186 patients. We refer to the combined and processed time-series data set as 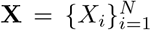 such that 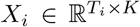 represents patient *i*’s *K*-dimensional time series of length *T*_*i*_ (i.e., each row is a visit at some time point, and 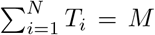. We associate **X** with the binary targets **Y** ∈ {0, 1}^*N*^ where 0 and 1 encode RRMS and SPMS labels, respectively. Conventionally since SPMS is a more advanced stage of MS which comes after RRMS, the positive class is set to SPMS.

**Table S3:**
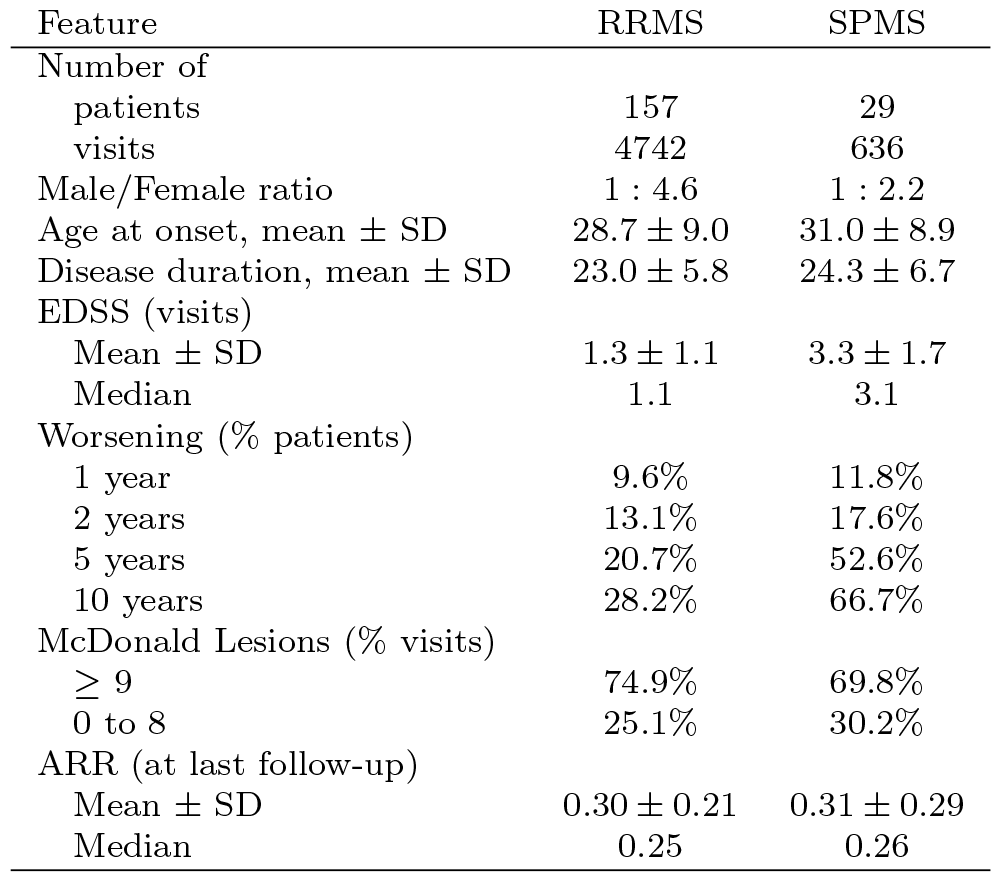
Summary characteristics of the RRMS and SPMS study cohorts.

**Fig. S2:**
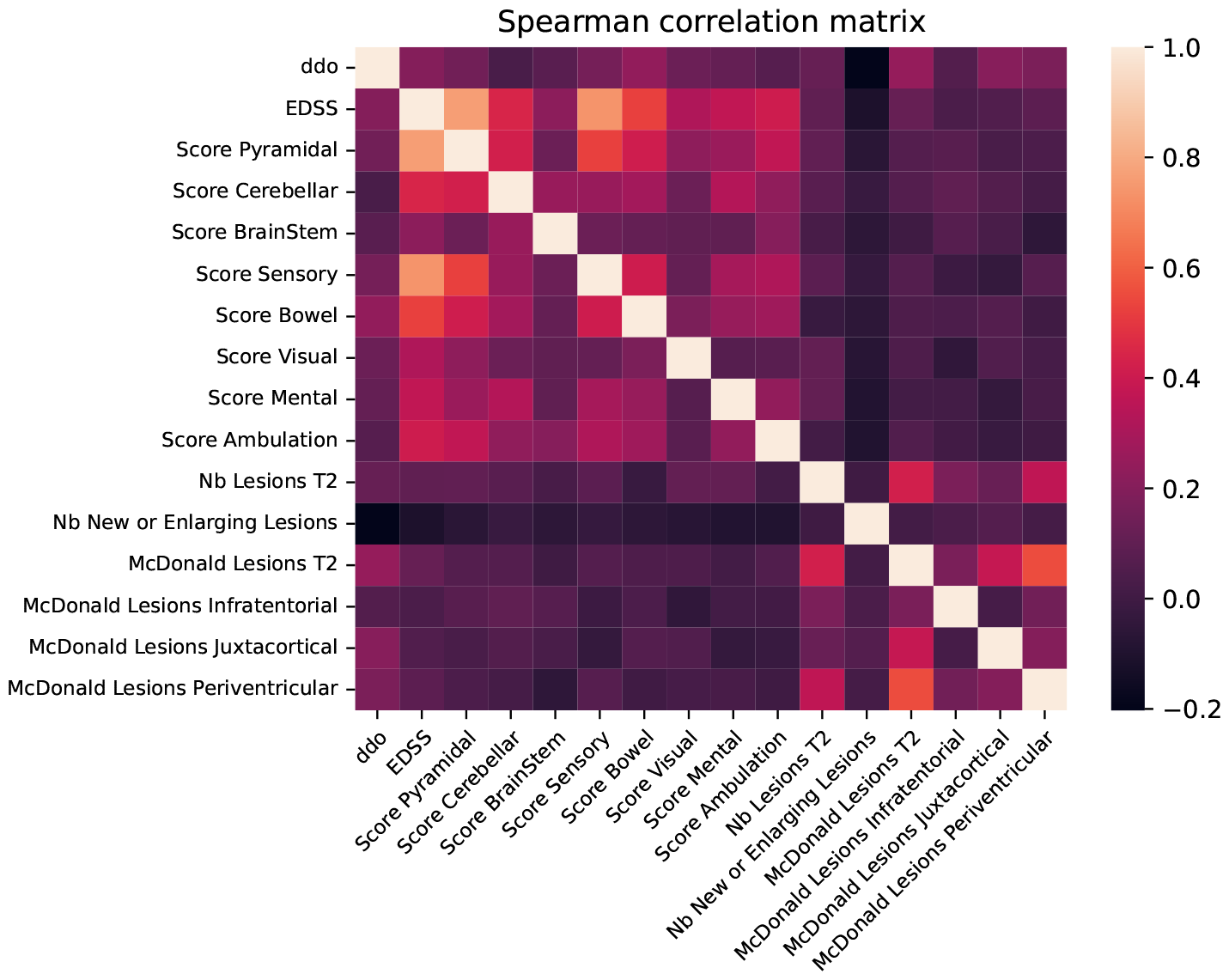
Heatmap of the Spearman correlations between input features.

### MS time-series data formatting

For the purpose of visualization, we seek a *d*-dimensional representation of the *K*-dimensional time series in 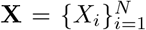 such that *d << K*. Usually, this is performed by a DR algorithm, but most are built to transform static data, which is not the case here. To solve this, we build the flattened data matrix **X**_*flat*_ ∈ ℝ^*M×K*^ which consists of the concatenation of all *X*_*i*_. Each row of the matrix **X**_*flat*_ corresponds to a *K*-dimensional visit for any patient. Thus, **X**_*flat*_ can be seen as the phase-space trajectory for the time-varying system determined by the patient time series in **X**. Its rows are also associated with **Y**_*flat*_ ∈ {0, 1}^*M*^ which links each visit to the outcome of the corresponding patient. Then, **X**_*flat*_ is passed through a given DR algorithm, jointly with **Y**_*flat*_ when supervised, producing a *d*-dimensional representation of patient visits from which *d*-dimensional patient time series are re-constructed thanks to the original patients’ identifiers and the timestamps of their visits.

Note that the above approach does not make explicit use of the longitudinal nature of the *N* observations. This is why we include time in the input feature set, and the longitudinal structure preservation of the original data set is taken into account in our evaluation framework. For the rest of our study, we refer to 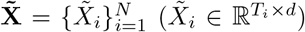 as the resulting *d*-dimensional counterpart of the original patient time series 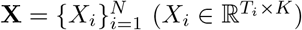.

**Fig. S3:**
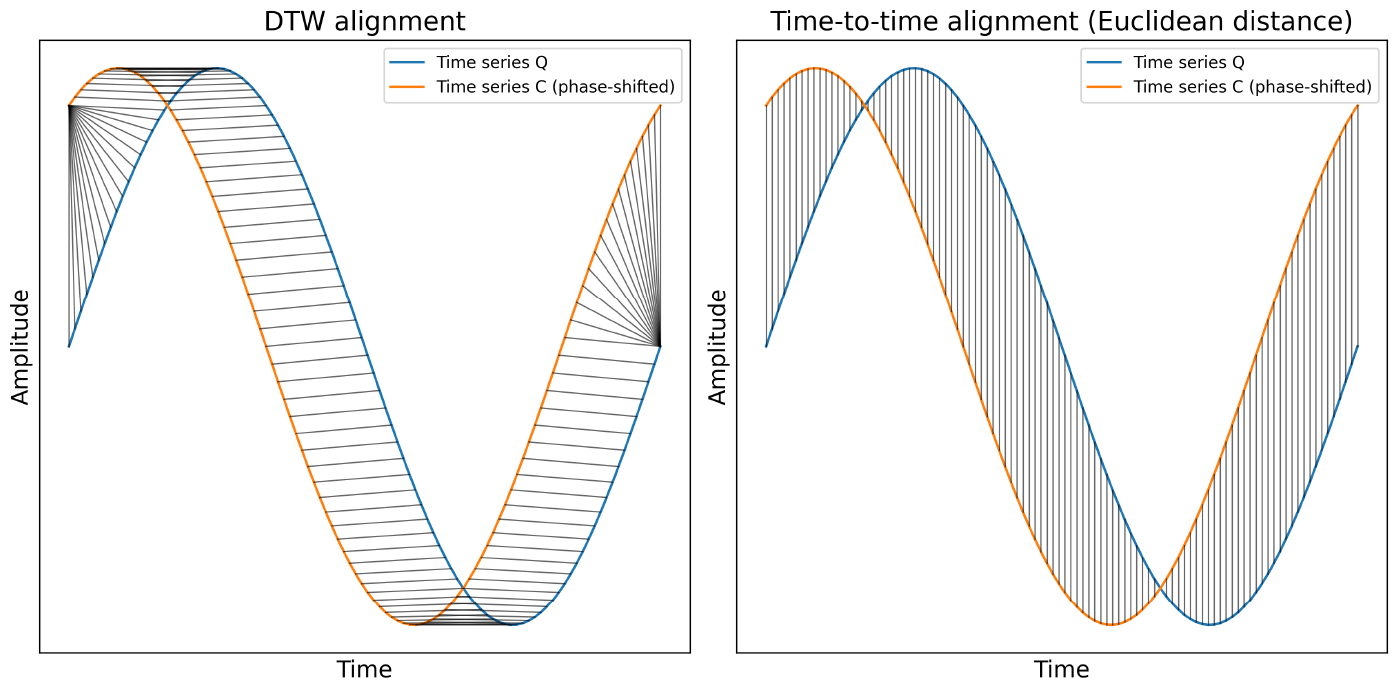
Comparison of DTW (left) and Euclidean (right) alignments between two time series of similar shape but not aligned in the time axis. The Euclidean distance sums the time-to-time distances between matching points, while DTW allows time distortions. With its restricted alignment, the Euclidean distance misses the similar shape shared by those two time series.

### Dynamic time warping

For time-series data, Euclidean distances are less meaningful and are thus replaced by dynamic time warping (DTW) [98, 99]. Given two time series *Q* and *C*, the DTW algorithm provides a dissimilarity measure *s*_*DT W*_ (*Q, C*) ≥ 0 based on a non-linear matching of respective points, as opposed to the Euclidean distance matching that calculates straight-line (geometric) distances (Fig. S3). More specifically, i f *n* a nd *m* d enote t he l engths o f *Q* a nd *C*, respectively, we write *Q* and *C* as *K*-dimensional ordered sequences,

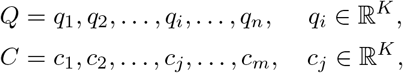

and construct an *n*-by-*m* matrix *M* such that each cell (denoted by their respective indices (*i, j*) for simplicity) contains the distance between the pair of points (*q*_*i*_, *c*_*j*_), usually the squared Euclidean distance: 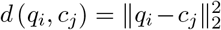. A warping path *W* is a set of distinct cells of the constructed matrix *M* that defines a mapping between *Q* and *C*. That is, 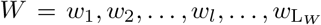 with *w*_*l*_ ∈ {1, …, *n*} × {1, …, *m*}. Then, we introduce the three following conditions to ensure that a warping path captures a “correct” sequence of (*q*_*i*_, *c*_*j*_) with respect to time, otherwise the time aspect of a time-series comparison would be completely lost:

i. boundary conditions: *w*_1_ = (1, 1) and 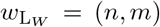, i.e., the warping path starts and ends in diagonally opposite corner cells of the matrix *M*. This prevents the occurrence of time warping being limited to a small section of the sequences;
ii. continuity: given *w*_*l*_ = (*a, b*), then *w*_*l−*1_ = (*a*^*′*^, *b*^*′*^), where *a* − *a*^*′*^≤ 1 and *b* − *b*^*′*^ ≤ 1. This restricts the allowable steps in the warping path to adjacent cells of *M* (including diagonally adjacent cells), as to avoid “time jumps”;
iii. monotonicity: given *w*_*l*_ = (*a, b*), then *w*_*l−*1_ = (*a*^*′*^, *b*^*′*^), where *a* − *a*^*′*^≥ 0 and *b* − *b*^*′*^≥ 0. This ensures that the warping path does not go backwards in time.

Finally, if **W** is the set of all possible warping paths subject to (i), (ii) and (iii), the DTW distance between *Q* and *C* is obtained from *W* ∈ **W** minimizing the warping cost 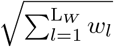

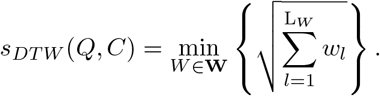

Under the above constraints (i), (ii) and (iii), DTW remains symmetric [100], that is, *s*_*DT W*_ (*Q, C*) = *s*_*DT W*_ (*C, Q*), so DTW provides an intuitive proximity between time-series objects and can be used similarly to Euclidean distance matching in a wide range of applications.

### Quantifying supervised embedding fit with DTW (MS Case)

The evaluation of the low-dimensional embeddings is inherently different for time-series data. Data points can no longer be viewed as static, independent observations. We want our embedding to capture the structure of the time series both locally and globally for meaningful exploratory analysis. We thus adapt our evaluation metrics LGS, LSP, and GSP to suit this purpose as follows.

**LGS**: In the context of our MS study, for the majority of cases where *i* and *j* represent different patients (*i* ≠ *j*), we have | *T*_*i*_ − *T*_*j*_ | *>>* 0 and patients’ time series are very likely to be recorded within different time (days from disease onset, (ddo)) periods. Moreover, physicians are rather interested in global long-term patterns of the course which is not restricted to time-to-time matching. This is why we make use of DTW which is more flexible than the traditional time-to-time Euclidean distance since it allows two time series that are similar but locally out of phase to align in a non-linear manner [101]. This variation of LGS still has the convenience of ranging between − 1 and 1 for easy interpretation.

**LSP**: The low-dimensional embedding should maintain the local variable importance structure for features useful for RRMS/SPMS classification. In our supervised context, features important to classify patients’ time series as RRM-S/SPMS should dominate the low-dimensional local structure of the embedded time series to be clinically insightful from a physician’s point of view. As in the non-time-series case, we produce two rankings of the input features with respect to their importance to the classification task and their importance to the local embedding structure generation. Clinically, important features can be thought of as the ones used by physicians in the retrospective RRMS and SPMS classification. However, our data also includes features whose importance is not *a priori* known, forcing us to objectively assess feature importance using a data-driven approach. To illustrate, we know that EDSS has been primarily used by physicians to diagnose the transition from RRMS to SPMS, but the impact of each of the 8 functional scores is hazy.

We numerically provide feature importance *𝒞* scores related to the RRMS/SPMS time-series classification by considering one feature at a time. First, we generate *K* data sets 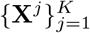 from **X** such that 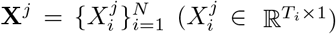 corresponds to the set of univariate patient time series determined by the *j*^*th*^ recorded feature. Then, for each **X**^*j*^ and the corresponding patient labels **Y**, we apply a leave-one-out cross-validation (LOOCV) to a DTW *k*-NN binary classifier such that the outputs are positive class probabilities. Since the numbers of RRMS and SPMS patients are respectively 157 and 29, the predicted probabilities are proportionally weighted to the inverse of the distances from the query points to mitigate class imbalance that inclines towards RRMS labels [102]. Finally, the importance score of feature *j* is computed as the area under the receiver operating characteristic curve (AUROC) based on the predicted class probabilities over the test folds of the LOOCV and their corresponding true classes. Thus, if *𝒞*^*j*^ denotes the resulting importance score of feature *j*, then the set of feature importances for the RRMS/SPMS classification task is defined as *𝒞* = {*𝒞*^*j*^ : *j* = 1, …, *K*}. Feature importances *ℒ* related to the local embedding structure generation are obtained similarly, except that we regress on the embedded time series, and each *ℒ*^*j*^ is determined by summing the DTW distances between the predicted time series and the true embedded time series across the test folds. As previously described, the local structure preservation score is defined as the correlation between the two sets of importance vectors, *𝒞* and *ℒ*.

**GSP**: We compute pairwise distances within each of the *K* univariate time-series data sets 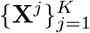 using DTW, and correlate each of the *K* corresponding condensed pairwise distances 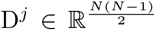 with the condensed pairwise distances (also using DTW) 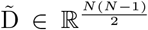 between the embedded patient time series 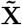. The score is then determined as previously described.

### Quantitative Experiment Details

Here we provide additional details about the quantitative comparisons of RFPHATE to other methods using our three metrics, LGS, LSP, GSP, in Tables 1 and 2.

For LSP and GSP, we used an unweighted *k*-NN model, setting *k* = 10 to determine feature importance using a permutation approach. The metric used to assess the goodness of fit of the *k*-NN models was overall accuracy for the label classification task, and *R*^2^ when regressing onto the embedding. 30 repetitions were used when extracting feature importance for stability. All other parameters were left as defaults. For these experiments, Spearman correlation was used for LSP and GSP, though Pearson’s correlation coefficient can also be used. We compared the performance of RF-PHATE against 23 other DR methods (10 unsupervised, 9 supervised, and 4 random forest-based methods [103]).

**Table S4:**
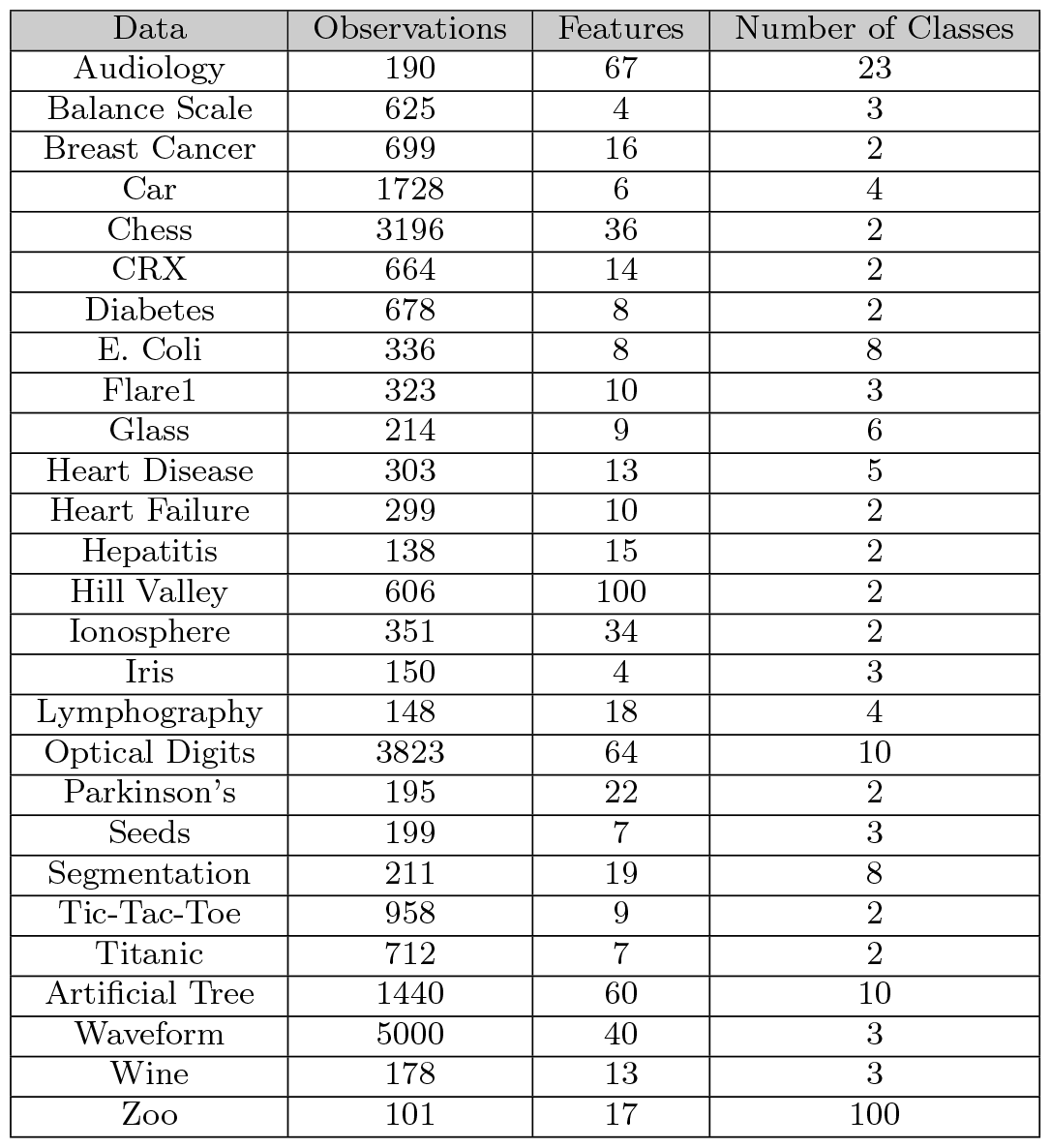
This table provides a brief summary of all datasets used to quantify the embedding fit using LGS, LSP, and GSP in Table 1. For each dataset, observations with missing values were removed.

| Data | Observations | Features | Number of Classes |
| --- | --- | --- | --- |
| Audiology | 190 | 67 | 23 |
| Balance Scale | 625 | 4 | 3 |
| Breast Cancer | 699 | 16 | 2 |
| Car | 1728 | 6 | 4 |
| Chess | 3196 | 36 | 2 |
| CRX | 664 | 14 | 2 |
| Diabetes | 678 | 8 | 2 |
| E. Coli | 336 | 8 | 8 |
| Flare1 | 323 | 10 | 3 |
| Glass | 214 | 9 | 6 |
| Heart Disease | 303 | 13 | 5 |
| Heart Failure | 299 | 10 | 2 |
| Hepatitis | 138 | 15 | 2 |
| Hill Valley | 606 | 100 | 2 |
| Ionosphere | 351 | 34 | 2 |
| Iris | 150 | 4 | 3 |
| Lymphography | 148 | 18 | 4 |
| Optical Digits | 3823 | 64 | 10 |
| Parkinson's | 195 | 22 | 2 |
| Seeds | 199 | 7 | 3 |
| Segmentation | 211 | 19 | 8 |
| Tic-Tac-Toe | 958 | 9 | 2 |
| Titanic | 712 | 7 | 2 |
| Artificial Tree | 1440 | 60 | 10 |
| Waveform | 5000 | 40 | 3 |
| Wine | 178 | 13 | 3 |
| Zoo | 101 | 17 | 100 |

We compared results across 27 publicly available datasets, with some details found in Table S4. No specific criteria were used to select the datasets; we tried to provide datasets with a variety of sizes and numbers of features. Across all datasets, the smallest number of observations was 98, and the largest was 5000. The number of classes ranged from 2 to 100. Due to the number of repetitions and number of embedding methods used we did not run this experiment with larger datasets. The number of features ranged from 4 to 100. Any observations with missing values were removed prior to embedding. All compared embeddings were two-dimensional. The number of remaining observations is reflected in the table. All continuous features were normalized to range from 0 to 1. Categorical features were encoded as integers.

For the MS quantitative analysis, we set the hyper-parameter *k* of our *k*-NN classifier/regressor to 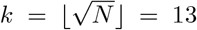. The metric used was DTW rather than Euclidean. Otherwise, default parameters were used. Classification importances *𝒞* are given in Table S5. The experiment was run 20 times and LGS, LSP, and GSP scores were each averaged across repetitions for each embedding. To qualitatively validate the quantitative comparison of the MS dataset, we also provide the 24 corresponding 3D plots of our MS database along with the ground truth univariate plot of historical EDSS in Fig. S4.

**Table S5:** Feature importances computed as the AUROC from LOOCV test predictions using a DTW *k*-NN classifier on univariate time series determined by each feature (see Methods), in decreasing order (higher is better). As expected, EDSS is the best predictor in the RRMS/SPMS classification.

| Feature name | AUROC |
| --- | --- |
| EDSS | 0.918 |
| Score Pyramidal | 0.877 |
| Score Ambulation | 0.829 |
| Score Cerebellar | 0.717 |
| Score Sensory | 0.706 |
| Score Bowel | 0.617 |
| Score Visual | 0.606 |
| Score Mental | 0.590 |
| Time (ddo) | 0.569 |
| Nb Lesions T2 | 0.568 |
| McDonald Lesions Juxtacortical | 0.543 |
| Score Brainstem | 0.541 |
| McDonald Lesions T2 | 0.511 |
| McDonald Lesions Infratentorial | 0.501 |
| Nb New or Enlarging Lesions | 0.495 |
| McDonald Lesions Periventricular | 0.394 |

### RRMS Cluster Details

Fig. 4a represents low-dimensional RF-PHATE patient trajectories with lines colored by their respective class labels. Given the high variability observed in patient courses within the RRMS group in Fig. 4a, we sought to determine more stable and typical groups of trajectories underlying RRMS by applying *k*-medoids clustering with *k* = 2 to the RRMS group. We used DTW as a proximity measure. Fig. 4b illustrates low-dimensional patient trajectories, with the RRMS patient trajectories colored based on the output of the clustering algorithm applied to the RF-PHATE embedding of the RRMS group of trajectories.

To facilitate a broader exploratory analysis that can be easily generalized, we chose *k* = 2 instead of higher values to maintain larger cluster sizes, resulting in 105 patients/3018 visits for the dark blue cluster, and 52 patients/1724 visits for the light blue cluster. To measure a level of spatial separation, we computed the silhouette score between these two clusters, which yielded a value of 0.59.

**Fig. S4:**
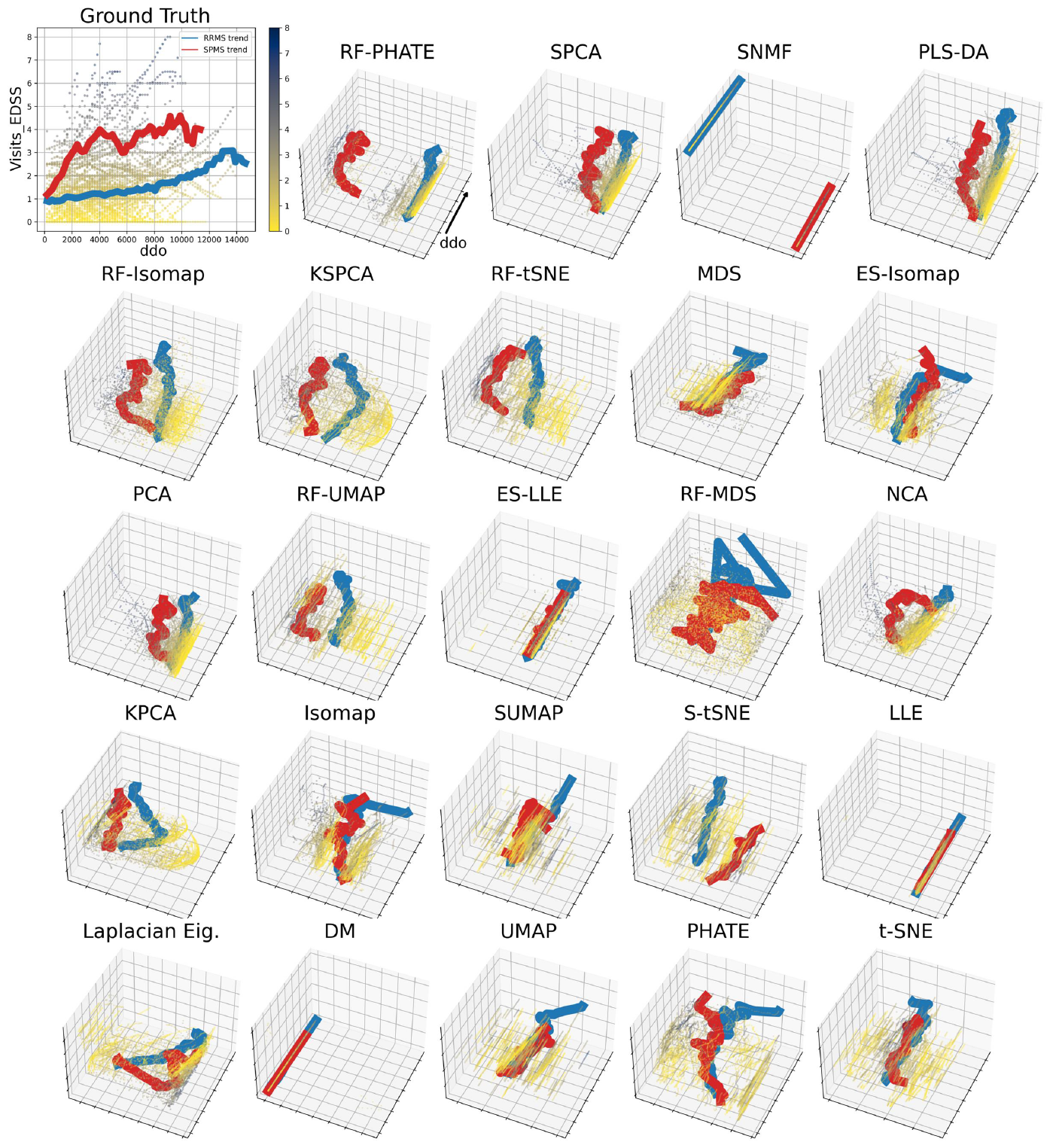
Three-dimensional visualizations of the MS time-series database using RFPHATE and 23 other embedding methods, sorted from left to right by their average rank in Table 2, along with the ground-truth EDSS patient trajectories (top-left plot). The three-dimensional plots have two axes for embedding dimensions and one for time, in days since MS onset (ddo). Each point represents a patient visit, colored by EDSS. Blue and red lines indicate the average trajectories of RRMS and SPMS groups, respectively. The plots align with our findings in Table 2. RF-PHATE remains the top choice for balancing class separability and structure preservation, while other embeddings perform progressively worse in both aspects as we move down the ranking.

### MS validation experiments

The MS validation database was collected by qualified neurologists from MS patients at the Neuro Rive-Sud clinic in Montreal. It consists of the same clinical features as in our original MS data. Owing to a significant shortage of imaging features, we excluded them from our input feature set. As indicated by their low contribution to the RF-PHATE embedding generation (Table S5), their removal did not undermine our validation experiment. We followed the same imputation and processing as in the initial MS database from the CRCHUM, resulting in 994 patients and 19,794 visits from which 10 features—EDSS, functional system scores and time, in days since disease onset— were recorded. Table S6 summarizes general characteristics of the validation data. Then, as in our initial experiments, we applied RFPHATE to the labelled visits, followed by a 2-medoids clustering on the RRMS group (Fig. S7a,b). Finally, we generated the same qualitative (Fig. S8) and quantitative (Fig S7c,d,e) plots to analyze and compare the three MS trends.

**Fig. S5:**
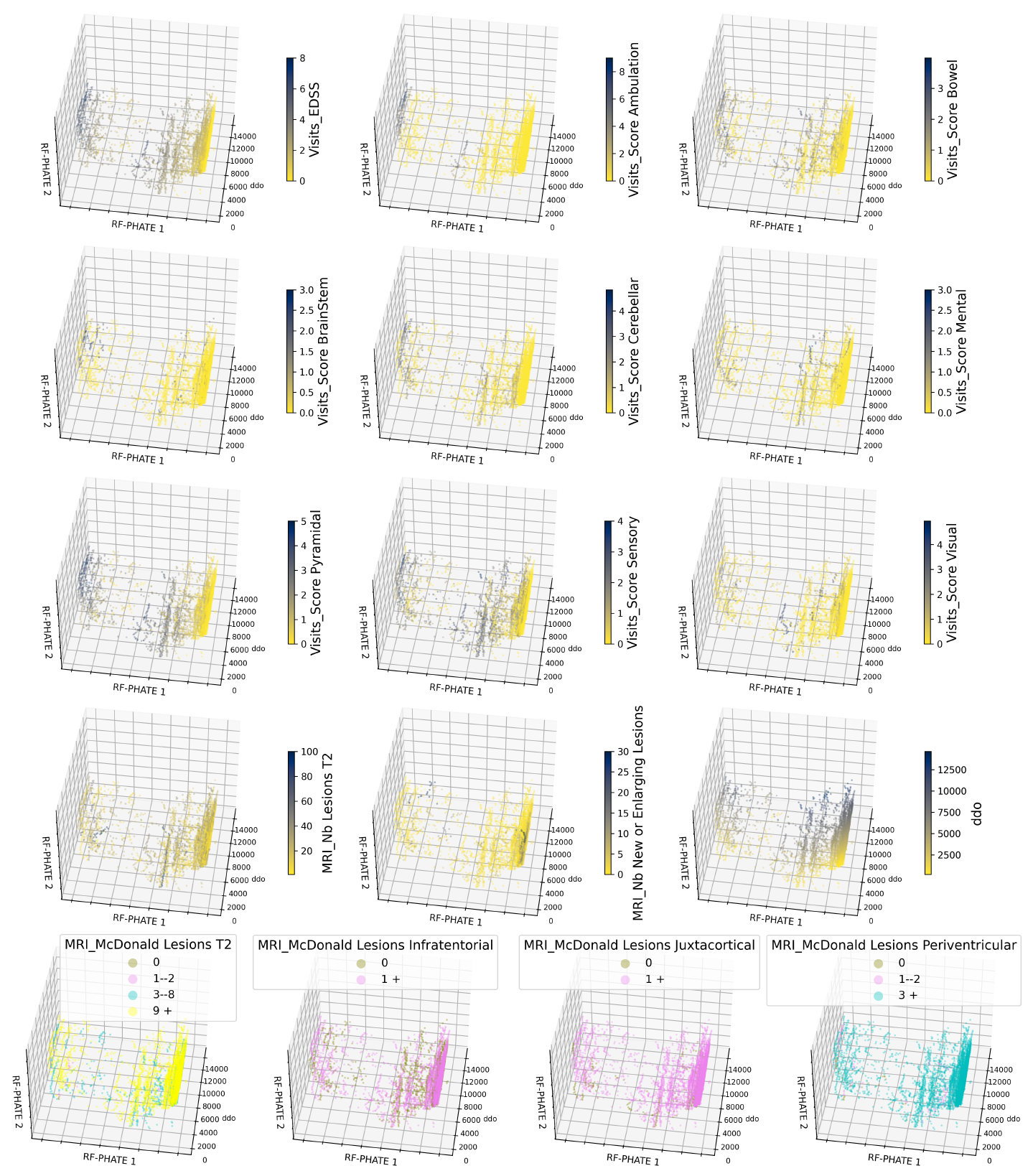
Three-dimensional RF-PHATE plots of MS patient visits colored by each of the 16 input features. The embedded patient visits within the RRMS group (right cluster) exhibit lower values compared to those in the SPMS group (left cluster) across nearly all clinical indicators. This observation is in line with the well-established disparities between RRMS and SPMS in terms of clinical worsening (domain knowledge retrieval). Simultaneously, we observe a substantial number of visits in the bottom-left region of the RRMS group with scores similar to those in the SPMS group. Still, some aspects, such as high ambulation scores, remain very specific to the SPMS group. These observations suggest the existence of a more severe subgroup of RRMS patients that was not explicitly labelled in our database (knowledge discovery). As expected, the plots related to MRI features do not highlight distinct patterns because the MRI features included in our input feature set hold limited importance in the RRMS/SPMS classification (Table S5).

**Fig. S6:**
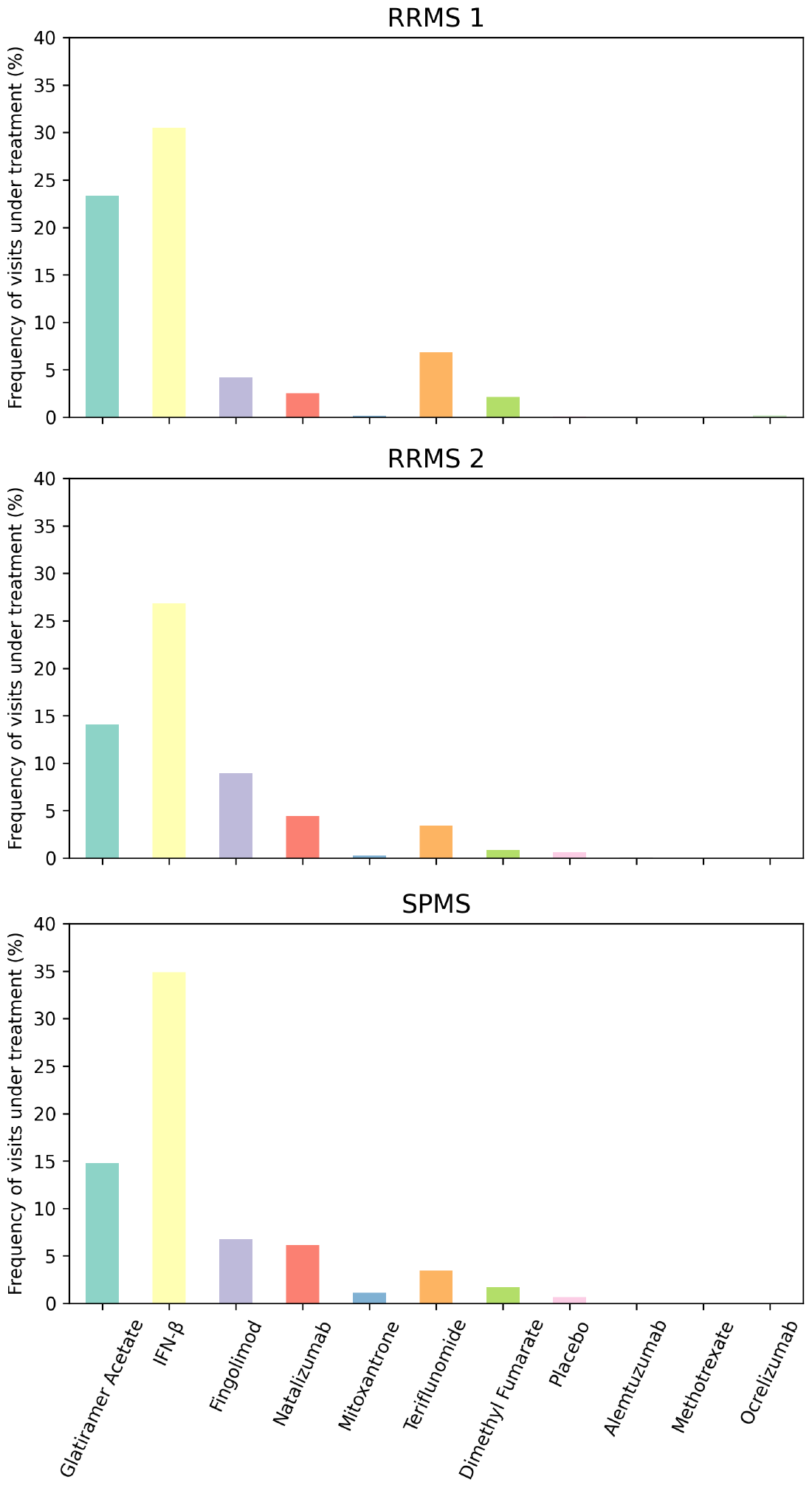
Frequency of patient visits under treatment with various MS-specific medications within the RRMS 1 (top), RRMS 2 (middle), and SPMS (bottom) clusters. Surprisingly, patients within the three clusters have been prescribed similar medications despite their distinct MS trajectories (Fig. 4b). This suggests that an incorrect treatment strategy may have been employed, as RRMS 2 and SPMS patients might have benefited from targeted treatments earlier to improve their outcome.

**Table S6:** Summary characteristics of the RRMS and SPMS validation cohorts.

| Feature | RRMS | SPMS |
| --- | --- | --- |
| Number of patients | 730 | 264 |
| visits | 14,387 | 5,407 |
| Male/Female ratio | 1 : 4.1 | 1 : 2.6 |
| Age at onset, mean ± SD | 34.0 ± 9.7 | 32.7 ± 10.2 |
| Disease duration, mean ± SD | 16.0 ± 9.0 | 27.1 ± 10.8 |
| EDSS (visits) |  |  |
| Mean ± SD | 1.9 ± 1.8 | 5.1 ± 2.0 |
| Median | 2.0 | 5.8 |
| Worsening (% patients) |  |  |
| 1 year | 8.6% | 6.8% |
| 2 years | 12.4% | 16.2% |
| 5 years | 21.3% | 42.0% |
| 10 years | 31.0% | 71.2% |
| ARR (at last follow-up) |  |  |
| Mean ± SD | 0.21 ± 0.24 | 0.15 ± 0.13 |
| Median | 0.15 | 0.11 |

### Raman data collection and processing

The A549 cells were purchased from ATCC (Manassas, VA, USA) and cultured in an F-12k medium containing 10% fetal bovine serum (Thermo Fisher Scientific, Waltham, MA, USA) at 37°C with 5% CO_2_ in a humidified atmosphere. DEPs (4mg/ml) were prepared with a culture medium, vortexed for 10 seconds, and then sonicated for 20 minutes at room temperature. The A549 cells underwent pretreatment in a plain medium, a medium containing resveratrol (RES), or one containing Cannabidiol (CBD) at 10 µM for 24 hours. Subsequently, cells were treated with a DEP solution at 0, 10, 25, 50, and 100 µg/ml for 24 hours.

The Raman spectra were measured by a Renishaw inVia Raman spectrometer (controlled by WiRE 3.4 software, Renishaw, UK) connected to a Leica microscope (Leica DMLM, Leica Microsystems, Buffalo Grove, IL, USA) equipped with a 785 nm near-infrared (IR) laser which was focused through a 63 ×NA = 0.90 water immersion objective (Leica Microsystems, USA). The standard calibration peak for the spectrometer with silicon mode at a static spectrum was 520.5 ± 0.1 cm^*−*1^. A549 cells were cultured on MgF_2_ (United Crystals Co., Port Washington, NY, USA) and imaged in Earle’s balanced salt solution (EBSS). Raman spectra between 600 and 1800 cm^*−*1^ wavenumbers were recorded for 1 accumulation of 10-second laser exposure in static mode. Five points on each cell were randomly selected for measurements. At least 75 spectra (five points for each cell and fifteen cells) in each condition were collected for further analysis. The raw Raman spectra were baseline corrected by adaptive iteratively re-weighted penalized least squares. The Python library called BaselineRemoval (version 0.1.4) was used [104]. The spectrum was further normalized to [0,1].

**Fig. S7:**
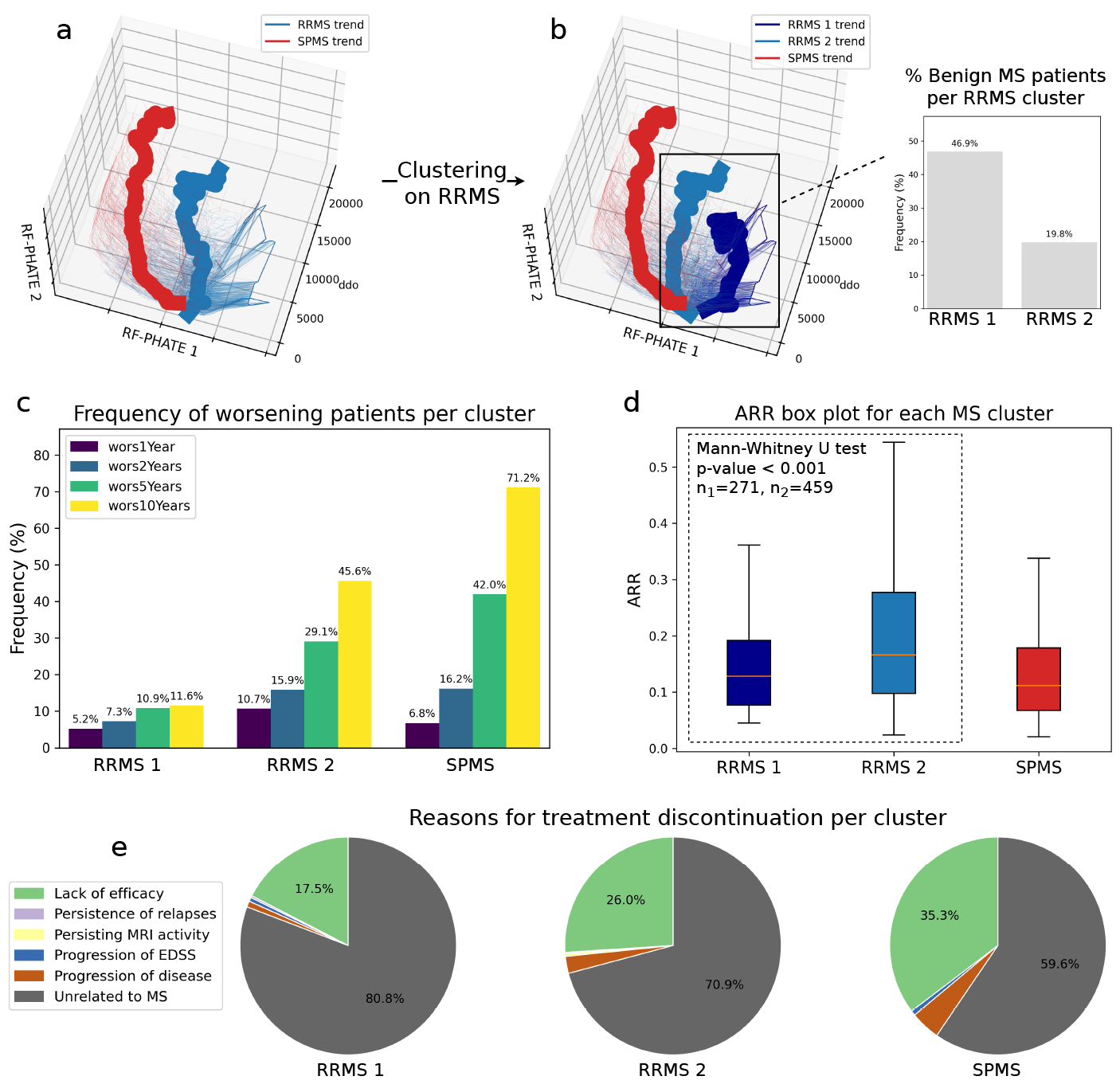
Exploratory analysis of our MS validation data using RF-PHATE. (**a,b,c,d,e**) Refer to Fig 4a,b,d,e,f for the legend. Overall, the same results hold for the validation data, confirming the existence of a severe relapsing-remitting subgroup of MS patients (RRMS 2).

**Fig. S8:**
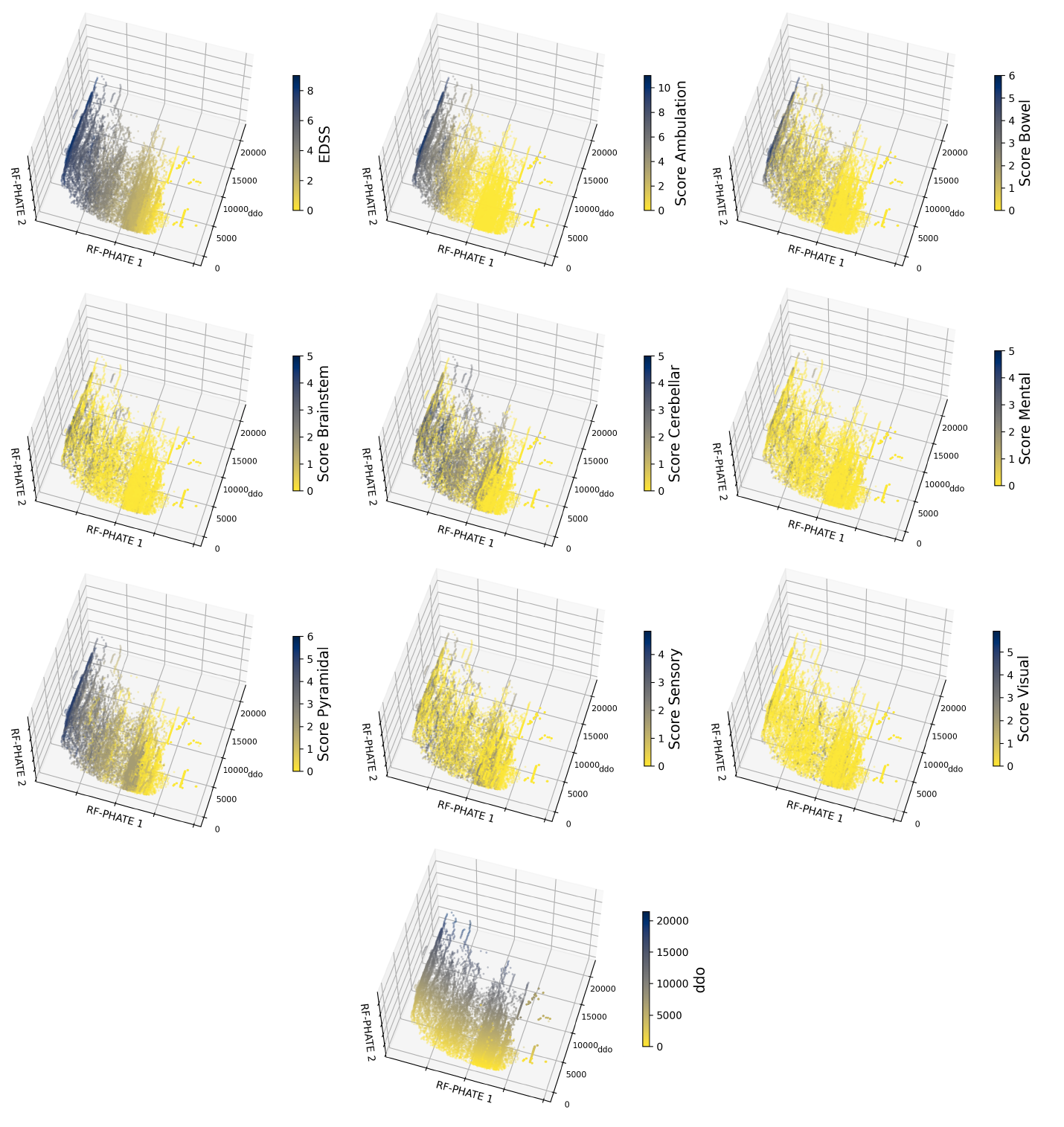
Three-dimensional RF-PHATE plots of MS patient visits from the validation data set, colored by each of the 10 input features. Visually, the trends seen in Fig. S5 remain consistent in these validation plots.

### Participants and samples – COVID-19 cohort

Plasma analytes were measured on prospectively enrolled COVID-19 individuals hospitalized between April 2020 and August 2021 with symptomatic infection with a positive SARS-CoV-2 nasopharyngeal swab (NSW) reverse-transcription polymerase chain reaction (RT-PCR) recruited into the Biobanque Québécoise de la COVID-19 (BQC19) [105]. Exclusion criteria were breakthrough or reinfection, plasma transfer therapy, or vaccination prior to infection. The study was approved by the respective IRBs (multicentric protocol: MP-02-2020-8929) and written informed consent was obtained from all participants or their legal guardians before enrollment. The research adhered to the standards indicated by the Declaration of Helsinki. COVID-19 hospitalized patients were stratified based on the severity of respiratory support at 11 days after symptom onset (DSO11): critical patients required mechanical ventilation (noninvasive ventilation, endotracheal intubation, extracorporeal membrane oxygenation), and non-critical patients required oxygen supplementation by nasal cannula or no supplemental oxygen. All-cause mortality was tracked up to 60 days after symptom onset (DSO60).

### Quantification of plasma analytes – COVID-19 cohort

Plasma vRNA, cytokine, and RBD-specific antibody concentrations were measured as previously described [57] at DSO11 (+/-4 days). Briefly, for SARS-CoV-2 viral RNA (vRNA), total RNA was extracted from 230 uL of plasma, and real-time PCR was performed using a N-specific primer of the SARS-CoV-2 genome. Purified RNA N transcripts (1328 bp) were quantified by Nanodrop (Thermo), and RNA copy numbers were calculated using END-MEMO. Cytokines were captured by magnetic beads array using a customized Human Luminex Discovery Assay (LXSAHM-26, R&D Systems) and acquired on the MagPix (Luminex). Only analytes significantly associated with a fatal outcome (p*<*0.01) in our previous work [58] were retained: TNFa, CXCL13, IL-6, IL-23, CXCL8, angiopoietin-2, RAGE, and Surfactant Protein D. RBD-specific IgG, IgM, and IgA were quantified using an in-house SARS-CoV-2 RBD ELISA assay, as described elsewhere [106]. Plasma from pre-pandemic uninfected donors was used as a negative control. The seropositivity threshold was established using the following formula: mean of all COVID-19 negative plasma + (3 standard deviations of the mean of all COVID-19 negative plasma).

### Data-processing - COVID-19 cohort

As in [57], we retain only plasma samples at DSO11 (+/-4 days) and if a patient has multiple samples in the considered time frame, we include only the one closest to DSO11. The final input consists of 242 plasma samples.

We use the same data processing as in [57] to run RF-PHATE, PHATE, and *k*-means. All 14 input features are log-scaled and standardized (0 mean, variance 1). For RF-PHATE, we set the diffusion parameter *t* to 12 and otherwise use the default parameters of the random forest estimator of scikit-learn v1.1.1.

### COVID-19 longitudinal data details

In this section, we provide additional details on the patient time series extracted from COVID-19 data [57]. The initial database contains 604 plasma samples from 304 patients. We followed the same data imputation and formatting as in our MS case study. Each patient time series consists of 14 immunological features recorded at different time points up to DSO30. We exclude five patients because one or more features are missing in their records.

We also added the time component (in days since disease onset) to the existing feature set, resulting in 1,425 plasma samples from 299 patients associated with 15 features. The label of interest is the all-cause mortality (Yes/No) of any patient 60 days after enrollment. The positive group, which includes deceased patients, consists of 40 patients and 256 samples, while the negative group, comprising non-deceased patients, consists of 259 patients and 1,169 samples. The data processing is otherwise identical to the COVID-19 cross-sectional data.

### Artificial tree construction

The artificial tree data used in Fig. 3 follows the construction provided in the original PHATE paper [1]. The tree is crafted as follows. The initial branch has 100 linearly-spaced points spanning four dimensions, with the rest of the dimensions set to zero. The second branch’s 100 points are constant in the first four dimensions, mirroring the first branch’s endpoint. The subsequent four dimensions progress linearly, while others are set to zero. Similarly, the third branch progresses in dimensions 9–12 instead of 5–8. Other branches follow suit but with varying lengths, adding 40 points at endpoints and branch points, and zero-mean Gaussian noise (s.d. 7). This models gene expression advancement along branches. Additional noise dimensions total the data’s dimensionality to 60. Prior to visualization, all features were normalized to fall between 0 and 1.

In the extremely noisy case (Fig. 3 with gray backgrounds), we added 500 variables simulated from a uniform (0, 1) distribution.

## Declarations

## Acknowledgments

We gratefully acknowledge MSBase Foundation for their collaboration in this research and the Canada CIFAR AI Chairs.

## Funding

This research was supported in part by a Natural Sciences and Engineering Research Council of Canada (NSERC) PGS D Scholarship [S.M.], Fonds de Recherche du Québec Nature et technologies (FRQNT) PhD Scholarship [S.M.], COVID-19 excellence scholarship from the Université de Montréal [E. B.-R.], American Foundation for AIDS Research (amfAR) grant 110068-68-RGCV [D.E.K., N.C., A.F.], Canada’s COVID-19 Immunity Task Force (CITF) in collaboration with the Canadian Institutes of Health Research (CIHR) grant VR2-173203 [D.E.K., A.F.], CIHR grant # 178344 [D.E.K., A.F.], National Institutes of Health (NIH) grant no. 1R15HL168697 [K.R.M.,A.Z.], Canada CIFAR AI Chair [G.W.], NSERC Discovery grant 03267 [G.W.], NIH grant R01GM135929 [G.W.], the T1 Canada Research Chair in MS [A.P.], the Canada Institute of Health Research [A.P.], MS Canada [A.P.], the NMSS [A.P.], and the Canadian Foundation for Innovation [A.P.].

The authors declare no competing interests.

## Ethics approval

MS data was collected from MS patients with full ethical approval (BH07.001, Nagano 20.332-YP). All of the MS validation data was collected at the Neuro Rive-Sud MS Clinic in Greenfield Park, Québec, Canada, affiliated with Charles LeMoyne Hospital, Faculty of Medicine, Sherbrooke University. The project was approved by the Charles LeMoyne Research Centre Ethics Committee.

## Code availability

A Python implementation of RF-PHATE is available on GitHub for academic use: https://github.com/jakerhodes/RF-PHATE

## Author contributions

K.R.M. and G.W. envisioned the project. J.S.R. implemented the method. A.C. assisted in the development of the method. J.S.R., A.A., S.Z., and S.M. performed the analyses. K.R.M., J.S.R., A.A., and S.M. wrote the paper. G.W., E.B.-R., D.E.K., S.Z., and A.P. assisted in writing. W.Z. and A.Z. provided the Raman data and assisted in the analyses. A.P., C.L., M.C., and S.Z. provided the MS clinical data from the CRCHUM. E.B.-R., A.Pa., L.M., N.C., A.F., D.E.K. provided the COVID-19 data. E.B.-R. compiled all lab and clinical COVID-19 data. F.G. provided the MS validation clinical data from the Neuro Rive-Sud clinic.

## Notes

https://github.com/jakerhodes/RF-PHATE

### Competing Interest Statement

The authors have declared no competing interest.

### Summary of Updates

Additional experimental data usage was added to the manuscript. Added new authors.

